# Intracranial Markers of Conscious Face Perception in Humans

**DOI:** 10.1101/037234

**Authors:** Fabiano Baroni, Jochem van Kempen, Hiroto Kawasaki, Christopher K. Kovach, Hiroyuki Oya, Matthew A. Howard, Ralph Adolphs, Naotsugu Tsuchiya

**Affiliations:** School of Psychological Sciences, Faculty of Biomedical and Psychological Sciences, Monash University, Australia; NeuroEngineering Laboratory, Department of Electrical & Electronic Engineering, The University of Melbourne, Australia; Institute of Neuroscience, Newcastle University, Newcastle upon Tyne, United Kingdom; Department of Neurosurgery, University of Iowa, Iowa City, IA, USA; Division of Humanities and Social Sciences, California Institute of Technology, Pasadena, CA, USA; Monash Institute of Cognitive and Clinical Neuroscience, Monash University, Australia; Decoding and Controlling Brain Information, Japan Science and Technology Agency, Chiyoda-ku, Tokyo, Japan

## Abstract

The comparison between perceived and unperceived trials at perceptual threshold isolates not only the core neuronal substrate of a particular conscious perception, but also aspects of brain activity that facilitate, hinder or tend to follow conscious perception. We take a step towards the resolution of these confounds by combining an analysis of ECoG neuronal responses observed during the presentation of faces partially masked by Continuous Flash Suppression, and those responses observed during the unmasked presentation of faces and other images in the same subjects. Neuronal activity in both the fusiform gyrus and the superior temporal sulcus discriminated seen vs. unseen faces in the masked paradigm and upright faces vs. other categories in the unmasked paradigm. However, only the former discriminated upright vs. inverted faces in the unmasked paradigm. Our results suggest a prominent role for the fusiform gyrus in the configural perception of faces.

## Introduction

In the last couple of decades, the relationships between brain activity and the contents of perceptual consciousness have been investigated using a variety of experimental techniques operating at different spatial and temporal scales, from single-unit, multi-unit and local field potential recordings in monkeys (Logothetis and Schall, 1989; Leopold and Logothetis, 1996; Wilke et al., 2006; Maier et al., 2007; Wilke et al., 2009), to non-invasive neuroimaging techniques such as EEG, MEG and fMRI in humans (e.g., (Tong et al., 1998; Grill-Spector et al., 2000; Dehaene et al., 2001; van Aalderen-Smeets et al., 2006; Liu et al., 2012; Schurger et al., 2015)) (see (Rees et al., 2002; Tononi and Koch, 2008; Dehaene and Changeux, 2011; Boly et al., 2013; Panagiotaropoulos et al., 2014) for reviews).

The scientific investigation of perceptual states presents a unique challenge, since it requires the objective measurement of subjective states. In particular, accuracy in reports of subjective states is a critical prerequisite for this investigation. With sufficient amount of training and careful experimental design (Leopold et al., 2003), monkeys (and potentially other animals) can be trained to report their perceptual states in a reliable manner (see, for example, (Leopold and Logothetis, 1996)). However, the investigation of the neuronal correlates of conscious awareness in human subjects constitutes a great advantage, since they can provide accurate reports of their perceptual states with minimal training following verbal instructions from the experimenter. This is critical, especially if graded levels of perceptual awareness are considered, as in the current study.

In humans, non-invasive neuronal recordings have been extensively employed in the search of the neuronal correlates of consciousness. Here, we recorded electrocorticography (ECoG) from subdural electrodes implanted on the ventral and lateral surface of the temporal lobes in five epileptic patients undergoing pre-surgical seizure monitoring while they engaged in visual perception tasks. Intracranial recordings from human subjects undergoing pre-surgical monitoring constitute a precious opportunity to advance our understanding of the neuronal correlates of conscious perception (e.g. (Kreiman et al., 2002; Gaillard et al., 2009; Fisch et al., 2009; Aru et al., 2012a; Willenbockel et al., 2012; Quiroga et al., 2014), see (Engel et al., 2005; Mukamel and Fried, 2012) for reviews), due to the direct measure of electrophysiological responses as well as their high spatial and temporal resolution in comparison with non-invasive modalities.

Several techniques have been proposed to investigate the neuronal correlates of conscious visual perception (Kim and Blake, 2005). These techniques enable the dissociation between retinal images and subjective perception.

Previous intracranial recording studies have investigated the neuronal correlates of conscious visual perception using stimuli that are perceptually degraded by a technique known as backward masking (BM) (Gaillard et al., 2009; Fisch et al., 2009; Quiroga et al., 2008). In a typical BM paradigm, a target image is presented briefly, followed by a masking image after a variable delay, known as Stimulus Onset Asynchrony (SOA). Short SOAs prevent the target image from being consciously perceived, while long SOAs allow the target image to emerge to consciousness reliably. At intermediate SOAs, conscious visibility fluctuates across trials. While most BM studies investigated the neuronal correlates of consciousness by comparing trials that differed markedly in either stimulus configuration (e.g., (Dehaene et al., 2001)) or other covariates, such as subject training (e.g., (Grill-Spector et al., 2000)), some recent studies aimed to more subtle contrasts that could more specifically expose the neuronal correlates of consciousness (Gaillard et al., 2009; Fisch et al., 2009). However, even these latter studies compared visible and invisible conditions in response to similar, but not identical, input stimuli, due to the experimental difficulty of adjusting SOA at perceptual threshold (but see (Quiroga et al., 2008; Del Cul et al., 2007) for examples where the contrast between seen and unseen targets at threshold SOA was possible for a subset of subjects). Thus, studies using BM may generally confound neuronal activity related to different perceptual outcome with neuronal activity related to different visual stimulation.

Here, we employed a different masking technique, known as Continuous Flash Suppression (CFS). This technique is based on the presentation of rapidly changing Mondrian patterns to one eye, while a static image (the target) is presented to the other eye ( (Tsuchiya and Koch, 2005), see (Yang et al., 2014; Sterzer et al., 2014) for recent reviews). Depending on the contrast of the target image and the Mondrian masks, the target image can be completely invisible, clearly visible, or visible only in a subset of the trials. The latter condition is of special interest, since the contrast between neuronal activity corresponding to trials with different visibility outcomes, in conditions of equal stimulus contrast, enables us to assess the neuronal correlates of visibility in the absence of any change in the physical properties of the stimulus.

Even when comparing trials corresponding to identical physical stimuli, but different perceptual outcomes, the resulting differences cannot be unambiguously considered as core neuronal correlates of phenomenal conscious perception, or NCC-core (Aru et al., 2012b; de Graaf et al., 2012; Miller, 2007; van Boxtel and Tsuchiya, 2015; Tsuchiya et al., 2015), because they likely reflect additional processes that also differ between the conditions. In fact, the comparison between seen and unseen trials can also reveal brain states that facilitate (e.g., attentional mechanisms) or hinder (e.g., mind-wandering) the perceptual awareness of threshold stimuli. In addition to this, conscious visual perception of target stimuli can trigger a cascade of neuronal processes related to memory formation, generation of associations and motor preparation for the ensuing response. Aru *et al.* and de Graaf *et al.* conveniently termed the potential confounds belonging to the former category as NCC-prs, or prerequisites, and the latter as NCC-cos, or consequences, of the conscious perceptual experience.

In this work, we take a step towards the dissociation between the neuronal correlates of core aspects of conscious visual experience (NCC-core) and their prerequisites and consequences by combining different, albeit related, experimental protocols. In particular, we considered a masked visual task, where stimuli were made partially invisible by Continuous Flash Suppression (CFS); and an unmasked visual task, where stimuli were clearly visible. Each of these tasks expose different cognitive processes: while the partially masked visual targets in the CFS task seemed to require some effort to be seen, unmasked images were clearly seen without effort.

Importantly, some of the stimuli used in the unmasked task (photographs of human faces) belong to the same category as the target stimuli in the masked task. Human faces constitute a stimulus category of exceptional behavioral and ecological relevance, and are known to be processed in specific circuits in ventral and lateral regions of the temporal lobe, most evidently in the Fusiform Gyrus (FG) and in the Superior Temporal Sulcus (STS) (Allison et al., 1994; Puce et al., 1995; Kanwisher et al., 1997; Haxby et al., 2000; Kanwisher and Yovel, 2006; Tsuchiya et al., 2008; Kawasaki et al., 2012).

Critically, the image categories used in the unmasked condition comprise inverted faces in addition to upright faces and non-face objects such as houses and tools. The comparison between neuronal activity in response to upright versus inverted faces is expected to reveal features of neuronal processing that are specific to configural or holistic perception, that is, a gestalt perception where the whole face is perceived in a qualitatively different manner from the sum of its parts (e.g., (Rossion and Gauthier, 2002)). This phenomenon can be measured behaviorally, for example via reaction times in recognition (e.g. same/different judgment) tasks.

The comparison between specific neuronal markers in the masked and unmasked conditions enables one to discard neuronal markers that could otherwise be considered as putative NCC-core if only the contrast between visible and invisible trials at threshold were considered. More generally, this work paves the way for a new promising set of methodological approaches in consciousness research based on the comparison between similar experimental protocols, which differ in specific aspects that expose the key differences that enable one to disentangle the different aspects of the conscious visual experience.

## Materials and Methods

### Data Acquisition

We recorded intracranially with electrocorticographic (ECoG) electrodes from 5 epilepsy patients undergoing pre-surgical monitoring. Sampling rate for the ECoG signal was 2034.5 Hz. Electrode location was based solely on clinical criteria. Patient age, gender, handedness, ocular and language dominance, and locations of seizure foci are reported in Table 1. We did not record for 12 hours after any generalized seizure event. The University of Iowa Institutional Review Board (IRB) approved the study (approval number 200112047), and written informed consent was obtained from each patient. Further details are reported in (Tsuchiya et al., 2008; Kawasaki et al., 2012).

**Table 1.**
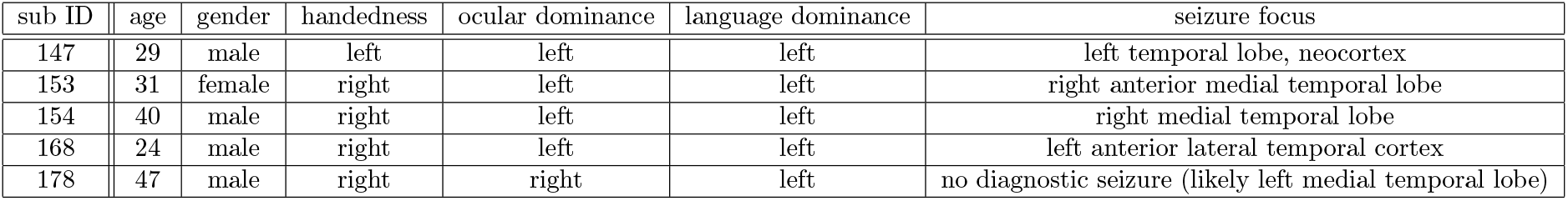
Demographic information for each subject.

| sub ID | age | gender | handedness | ocular dominance | language dominance | seizure focus |
| --- | --- | --- | --- | --- | --- | --- |
| 147 | 29 | male | left | left | left | left temporal lobe, neocortex |
| 153 | 31 | female | right | left | left | right anterior medial temporal lobe |
| 154 | 40 | male | right | left | left | right medial temporal lobe |
| 168 | 24 | male | right | left | left | left anterior lateral temporal cortex |
| 178 | 47 | male | right | right | left | no diagnostic seizure (likely left medial temporal lobe) |

### Electrode Localization

For each subject, we obtained structural T1-weighted MRI volumes (pre-and post-electrode implantation), CT scans (post-implantation) and digital photos of the electrodes (during surgery, only for the lateral temporal grid electrodes). Coronal MRI slices were obtained with 1 mm slice thickness, 0.78 x 0.78 mm in-plane resolution. Axial slices of the CT scans were obtained with 1 mm slice thickness, 0.45 x 0.45 mm in-plane resolution. Post-implantation CT scans and pre-implantation MRI were rendered into 3D volumes and co-registered using AFNI (NIMH, Bethesda, MD, USA) and/or ANALYZE software (version 7.0, AnalyzeDirect, KS, USA) with mutual information maximization. The resulting electrode locations for each subject are shown in Fig. 4A and S4.

### Behavioral Tasks

In order to assess and characterize neuronal activity related to conscious visual perception, we used two different sets of tasks: one involving masked images of faces, and another involving unmasked images of faces and other objects. The masked and unmasked tasks differ in cognitive requirements such as those related to attention, memory and report; hence their combined analysis can more specifically highlight neuronal activity directly related to the core mechanisms of conscious vision than could be possible if only the unmasked task were considered. In both sets of tasks, images were presented at fixation on a 19” ViewSonic VX922 LCD display (refresh rate: 60 Hz) and subtended about 7.5 x 10 deg in visual angle. Behavioral responses were collected using key presses on a USB keypad. We presented the stimuli using Psychtoolbox (Brainard, 1997) version 2.54 and MATLAB version 7.8 or higher on a PC running Windows XP. In order to ensure maximal precision in the temporal alignment of neuronal signals and visual stimuli, we displayed a small rectangle on the top-left corner of the screen, which changed in luminance in synchrony with the stimuli displayed at fixation, and recorded the response of a photodiode directly attached at that corner. The output from the photodiode was recorded along with the electrophysiological responses in the same recording system and used for segmenting the raw ECoG traces (see subsection “Data Analysis”).

#### Masked Visual Task (CFS)

In each trial, subjects were presented with a fixation cross displayed at the center of the screen and initiated a trial by pressing the space bar. The beginning of a trial was reflected on the screen by a 45-degree rotation of the fixation cross. Each trial consisted of two 200 ms intervals. After a variable period (uniformly distributed in [500,700] ms), the first interval was presented on the screen. Subsequently, after a variable period (uniformly distributed in [900,1100] ms), the second interval was presented on the screen.

In both intervals, three distinct Mondrian patterns were flashed within a frame composed of black and white squares to the dominant eye to suppress a visual input to the non-dominant eye (Continuous Flash Suppression (Tsuchiya and Koch, 2005)). Each Mondrian pattern was presented for 67ms and updated without any blank between the different patterns. In one of the two intervals, a face image was presented to the non-dominant eye, while in the other interval a blank field was presented instead within the corresponding area of the black and white frame (Fig. 1A).

**Figure 1.**
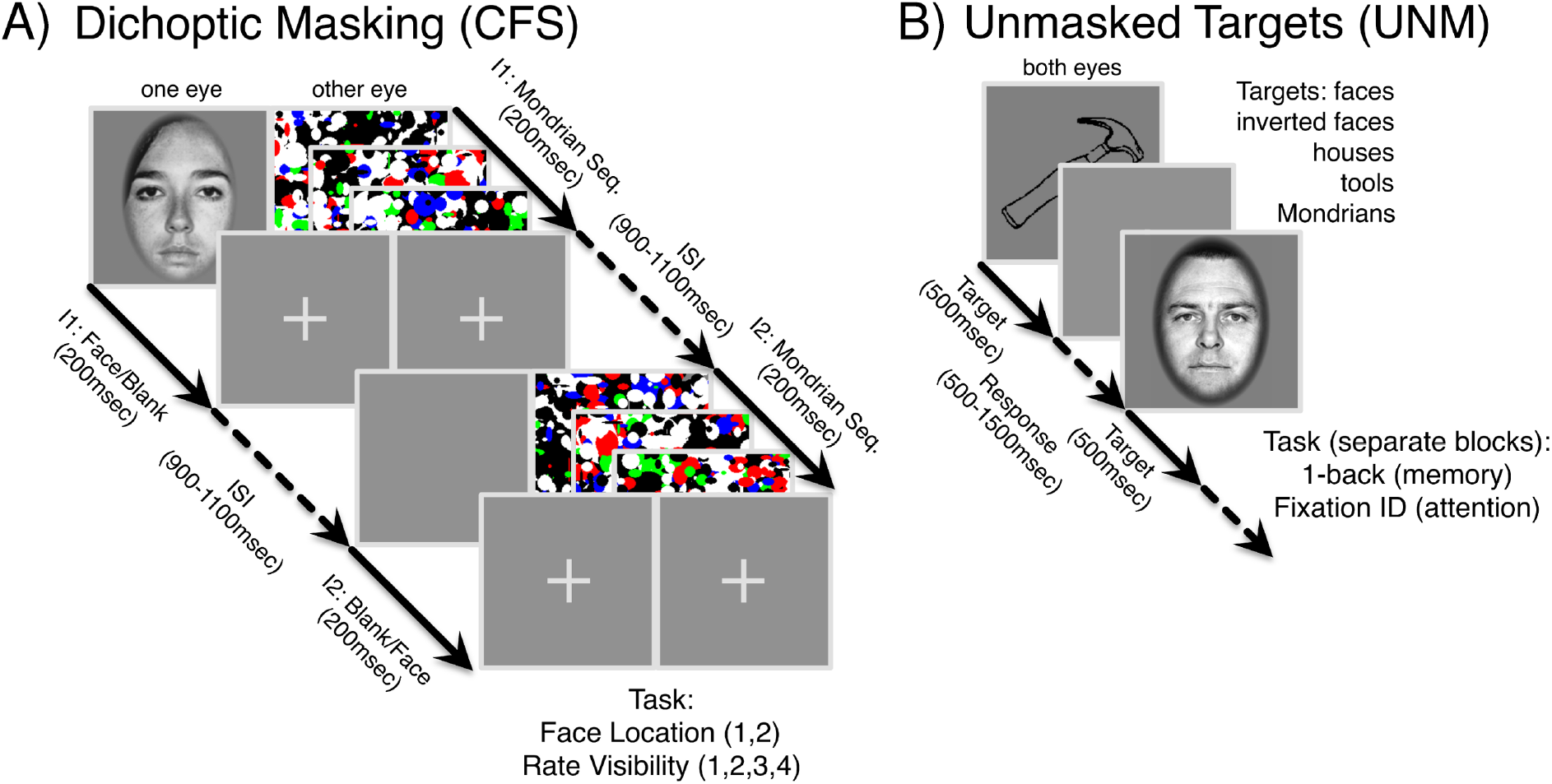
Experimental Protocols. (A) Masked condition (CFS). (B) Unmasked condition.

Following the termination of the second interval, after a variable period (uniformly distributed in [900,1100] ms), a first response screen appeared, asking for the interval which included the face (two-Interval Forced Choice, 2IFC). After the subject response, a second response screen appeared, asking for a face visibility rating, expressed according to the Perceptual Awareness Scale (PAS, (Overgaard et al., 2006), four-Alternative Forced Choice). PAS measures the subjective awareness of having seen a face, and ranges from 1 (the face has not been seen) to 4 (the face has been seen clearly).

While it is possible that slightly different results could have been obtained with confidence judgments (that is, judgments on the 2IFC task performance (Sandberg et al., 2010; Overgaard and Sandberg, 2012; King and Dehaene, 2014)), visibility ratings are more directly relevant to our primary concern (i.e., conscious face perception) than confidence judgments. Moreover, they have been shown to correlate more closely with objective performance and to yield a lower unconscious performance in identification tasks (Sandberg et al., 2010; Overgaard and Sandberg, 2012), suggesting that they might be generally more trustworthy than confidence judgments.

We used four different face identities with either neutral or fearful emotional expression (Ekman and Friesen, 1976) to reduce low-level perceptual learning (e.g., (Fahle, 2009)). Face images were presented at three logarithmically spaced contrast levels (with the exception of subject 178, for whom a different set of contrast values was used). Ideally, low contrast trials (*c*=1) would result in an objective performance in the 2IFC task near chance level, high contrast trials (*c*=3) would result in an objective performance around 90% or above, while intermediate contrast trials (*c*=2) would result in an objective performance around 75%. However, time constrains due to clinical requirements did not enable us to adjust contrast levels individually for each subject. Hence, a set of contrast levels were used for the first session. If the behavioral performance was too high or too low (e.g., objective performance above 85% or at chance level at intermediate contrast), contrast levels were scaled geometrically in successive sessions.

#### Unmasked Visual Task

Subjects were presented with images from different categories, presented at fixation for a duration of 500 ms (Fig. 1B). The stimuli used in each experiment, the behavioral response required and the number of sessions and trials for each condition are reported in Table 2.

**Table 2.** Stimuli, tasks and number of trials in each class for each subject in the unmasked condition. Stimulus set A comprises 15 images per category for upright faces, inverted faces, and houses; 50 images for tools; 100 for Mondrian patterns. Set B comprises 15 natural images for each category: faces (including upper half body), animals, landmarks, vehicles, flowers. Tasks: CD, Change Detection task on the fixation cross; OB, One-Back task for the stimulus category. In each trial, subjects report a change in the fixation cross (in the CD task) or a repetition of stimulus category (in the OB task) in a time window of duration 0.5 s or 1 s, respectively, immediately following stimulus offset. Following the termination of the response period, after a variable delay (uniformly distributed in [0,500] ms), the next trial began. ^1^ : this subject performed gender and emotion discrimination tasks as described in (Tsuchiya et al., 2008); the number of trials indicated refer to a “face” vs. “checkerboard” decoding analysis. In the case of subject 154, the set of recorded electrodes differed across sessions: “154a” and “154b” indicate the sets of electrodes that were recorded in only a subset of the sessions (shown in Fig. S4), while “154” indicates the set of electrodes that were recorded in every session.

| sub ID | Stimuli | # sessions (CD, OB) | U Face vs. Other | U vs. I Face |
| --- | --- | --- | --- | --- |
| 147<sup>1</sup> | Face, Checkerboard | 2 | 320/80 |  |
| 153 | Set A | 2,2 | 118/455 | 118/113 |
| 154a | Set A | 1,1 | 61/239 | 61/56 |
| 154b | Set A | 1,1 | 64/236 | 64/57 |
| 154 | Set A | 2,2 | 125/475 | 125/113 |
| 168 | Set A | 2,0 | 41/159 | 41/43 |
| 178 | Set B | 2,2 | 100/400 |  |

### Data Analysis

#### Behavioral Analysis

For each subject and face contrast value, we calculated the objective performance (defined as the ratio of the number of correct trials over the total number of trials) in the 2IFC task and counted the number of trials corresponding to each visibility rating. As expected, increasing face contrast values generally corresponds to an improvement in both objective performance and subjective visibility rating (Fig. 2A-B). Please note that the physical face contrast values used differ across subjects, since we aim to an objective performance around 75% at the intermediate contrast level for each subject.

**Figure 2.**
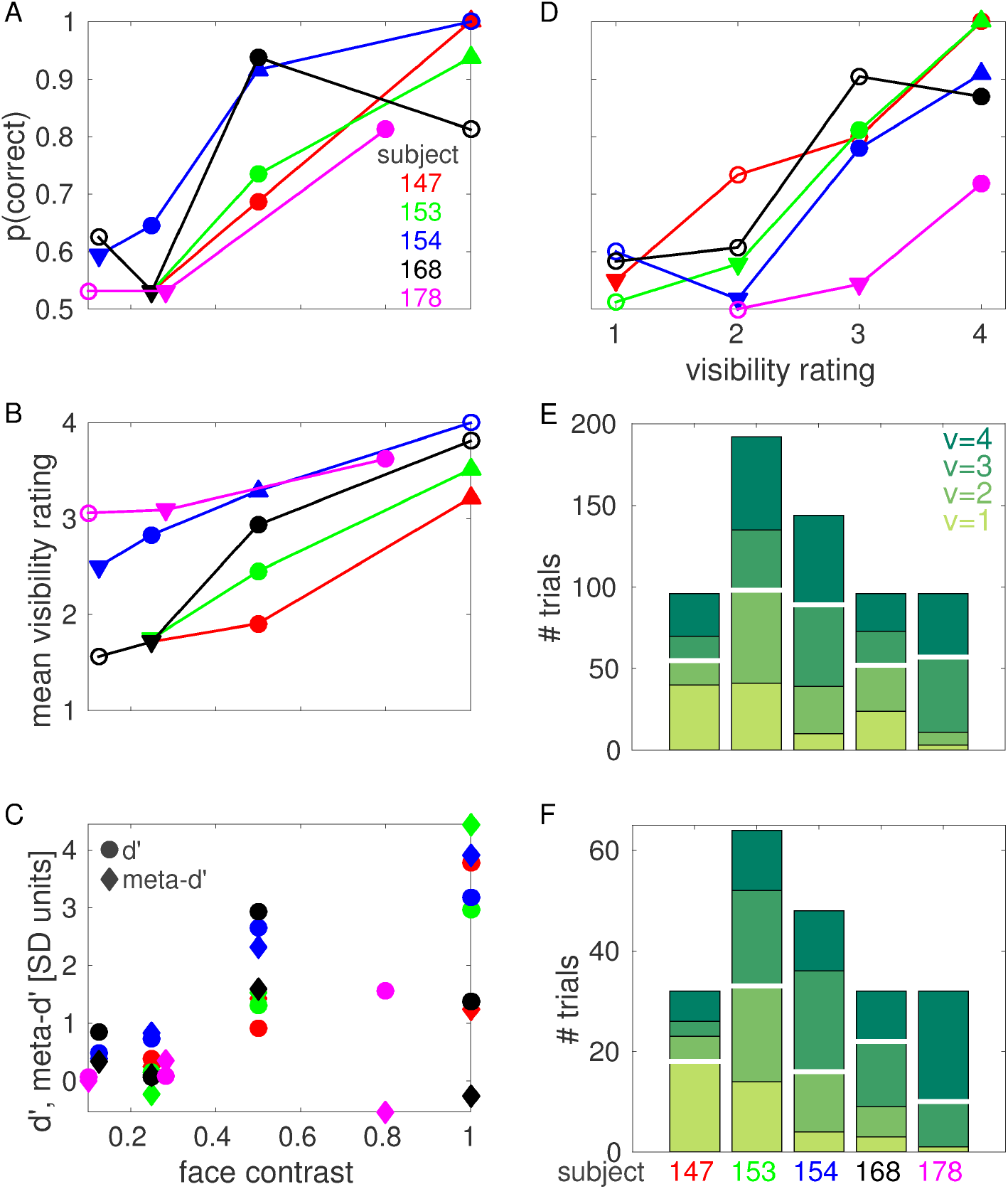
CFS behavioral results. (A) Objective performance (probability of correct response) at each face contrast value for each subject. (B) Mean visibility rating at each face contrast value for each subject. In A and B, filled circles indicate threshold face contrast *c_thr_*, downward triangles indicate low contrast *c_L_*, upward triangles indicate high contrast *c_H_*. Downward triangles indicating cL coincide for subjects 147, 153 and 168 in panel A. (C) d’ (circles) and meta-d’ (diamonds) at each face contrast value for each subject. (D) Objective performance at each visibility rating for each subject. For each subject, only visibility ratings that were reported in at least 8 trials are shown. Filled circles indicate threshold visibility rating *v_thr_*, downward triangles indicate low visibility *v_L_*, upward triangles indicate high visibility *v_H_*. (E) Number of trials for each visibility rating for each subject. (F) Number of trials for each visibility rating at threshold face contrast for each subject. The number of trials for each visibility rating at each face contrast value for each subject is reported in Fig. S1. White horizontal lines in E, F indicate median splits.

We measured the degree to which visibility ratings were predictive of objective performance (i.e., metacognition, or the ability to introspect on the accuracy of one’s own perceptual judgements) using a recently introduced measure from signal detection theory known as meta-d’ (Maniscalco and Lau, 2012; Barrett et al., 2013). In most cases, meta-d’ have the same sign and, often, similar amplitude as d’, indicating that high visibility ratings are predictive of correct objective performance, and vice versa, to an extent which is compatible with objective performance and visibility rating originating from a common (or largely redundant) internal signal (Fig. 2C, see also Fig. 2D for a similar analysis performed collapsing across contrast values). This relationship between visibility rating and performance justifies our treatment of visibility ratings as useful representations of subjects’ perceptual states. We note, however, that this relationship did not hold for subjects 168 and 178 at the highest contrast value we investigated, suggesting the possibility of a metacognitive impairment or poor understanding/execution of the task for these subjects (see also Discussion).

In this work, we aim to assess the neuronal correlates of subjective conscious perception, hence we compared trials that resulted in different perceptual outcome as reported by subjects, i.e. different visibility rating. In order to increase sample size, we grouped trials into a high visibility and a low visibility class using a median split of the data (Fig. 2E,F). The median split was determined independently for each classification considered, for each subject and, in the case of one subject where different electrodes have different numbers of trials (subject 154), for each electrode. The numbers of trials for each visibility rating for subject 154 indicated in Fig. 2E,F (and the corresponding median splits) correspond to those electrodes that were recorded in every session (i.e. those electrodes with the maximum number of trials for this subject). The number of trials for each visibility rating for the sets of electrodes that were recorded in only a subset of the sessions are reported in Fig. S1.

In order to investigate the neuronal correlates of conscious visual awareness in the absence of changes in physical properties of the presented stimuli, we compared trials with the same face contrast value but different visibility rating. This analysis was conducted using face contrast values that resulted in the face image being detected in roughly 75% of the trials. As is clear from Fig. 2B,D, different subjects adopted different criteria when declaring their degree of perceptual awareness: some subjects responded with high visibility ratings even when performing the task at chance level (e.g., subject 178), while others were much more conservative and responded with low visibility ratings even when performing the task with high accuracy (e.g., subject 147). To compensate for individual biases in visibility ratings, we adopted a definition of “threshold face contrast” based on the objective performance in the 2IFC task: threshold face contrast *c*_thr_ was defined for each subject as the lowest contrast value investigated that resulted in objective performance above 64% (indicated with filled circles in Fig. 2A; mean across subjects: 76%; range: 65% - 94%). We also considered a low contrast *c*_L_, defined as the highest contrast lower than *c*_thr_ that resulted in objective performance below 60% (mean across subjects: 54%; range: 53% - 59%); and a high contrast *c*_H_, defined as the lowest contrast higher than *c*_thr_ that resulted in objective performance above 90% (this condition was only realized in three subjects; mean across subjects: 95%; range: 92% - 100%).

Analogously, we investigated the neuronal correlates of changes in physical contrast in the absence of changes in the reported visibility rating. To this end, we compared trials with the same visibility rating but different face contrast. This analysis was conducted using trials with “threshold visibility rating” v_thr_, defined for each subject as the lowest rating with sufficient number of trials (see subsection “ECoG decoding analyses” for details) that resulted in objective performance above 64% (only realized in four subjects; indicated with filled circles in Fig. 2D; mean across subjects: 79%; range: 72% - 87%). We also considered a low visibility rating v_L_, defined as the highest rating lower than v_thr_ that resulted in objective performance below 60% (only realized in four subjects; mean across subjects: 55%; range: 52% - 58%); and a high visibility rating v_H_, defined as the lowest rating higher than v_thr_ that resulted in objective performance above 90% (only realized in two subjects; mean across subjects: 95%; range: 91% 100%).

#### Data preprocessing

ECoG signals were recorded with reference to the electrode placed under the scalp near the vertex of the skull. We bipolar re-referenced the original signals along the vertical and horizontal directions to remove low spatial frequency components and hence obtain a more localized signal and better exploit the fine spatial resolution of ECoG recordings. We removed 60 Hz line noise from the photodiode trace using a linear combination of sinusoids estimated using the MATLAB function rmlinesmovingwinc.m (included in the Chronux data analysis toolbox (Mitra and Bokil, 2007), http://chronux.org). Onset times for the two intervals in each trial were estimated as threshold-crossing times of the de-noised photodiode traces. Then, the onset times were used to segment the data in time windows comprising [-500,1500] ms relative to interval onset.

#### ECoG spectrogram analysis

We used the Chronux data analysis toolbox to estimate the spectrograms of the bipolar ECoG signals using a multi-taper method (Mitra and Bokil, 2007). We used 3 tapers and a time window of 100 ms (which corresponds to a half bandwidth of 20 Hz), slided in steps of 50 ms. To improve visualization and yield a distribution that is closer to normal, the logarithm of the power spectrum was considered for plotting and subsequent analyses. Other transformations (e.g., cubic root) were also considered and yielded comparable results.

#### ECoG decoding analyses

We estimated the amount of information conveyed by neuronal signals using binary Regularized Least-Square Classifiers (RLSC, (Rifkin et al., 2003)) with regularization parameter λ = 10^6^. Regularized Least-Square Classification is a machine learning technique that estimates the linear separability between patterns according to their class. Here, we aim to assess the amount of information conveyed by a spectro-temporal representation of the ECoG signal in each trial about the presented physical stimulus or the reported phenomenal experience. In particular, we considered log power at 10x11 (time,frequency) points for each trial, sampled from a uniform grid in the interval [100,600] ms after stimulus onset x [0,200] Hz, as the input to the classifiers (Tsuchiya et al., 2008).

A set of weights that optimally separate trials according to their class is determined using a subset of the available trials, denoted as training set. The performance of the classifier is defined using a different set of trials, denoted as test set, as the area under the Receiver Operating Characteristic (ROC) curve, which we refer to as A’ (A prime). We report the average A’ values over *N*_iter_ cross-validations. In each cross-validation, we randomly chose a set of 0.7 x min(*N*_1_, *N*_2_) (rounded to the nearest integer) trials of each class as the training set, where *N*_1_ and *N*_2_ are the number of trials in class 1 and 2, respectively. As the test set, we chose min(*N*_1_, *N*_2_) — round(0.7 x min(*N*_1_, *N*_2_)) trials of each class among those that are not already included in the training set. Before being fed to the classifier, inputs were z-transformed: the mean and standard deviation of log power at each time-frequency point in the training set was calculated, and used to transform both training and test sets. Then, optimal RLSC weights were estimated using training trials, and their capacity to separate test trials according to their class was measured as the area under the ROC curve (A’). The number of cross-validations *N*_iter_ was set to 100 for all the decoding analyses, except *v@c* and *c@v* decoding analyses, where *N*_iter_ = 1000 was used in order to decrease the greater sampling variability that results from decoding analyses on smaller samples.

Significance of A’ values was estimated via a permutation-based technique. For each classification considered, the class labels were randomly shuffled. Then, the average A’ value over *N_iter_* realizations of training and test sets was calculated as described above. This procedure was repeated *N*_perm_=1000 times, yielding a probability distribution of average A’ values corresponding to the null hypothesis of lack of linear separability between the two classes. An empirical average A’ value was considered significant at level p if it exceeded the p-percentile of the corresponding null distribution (p=0.05, p=0.01, p=0.001).

Significance thresholds at p=0.05 and p=0.01 were estimated separately for each classification considered. In order to improve the estimation of the significance threshold at p=0.001, null A’ values were pooled across electrodes, and the corresponding significance threshold was calculated from the resulting null distribution. For each subject and electrode, each analysis was only considered if at least 10 trials were available in the least populated class.

#### Face responsiveness

Our purpose is to identify brain loci that are part of the neuronal correlates of conscious face perception, hence we restricted our analysis to electrodes that are responsive to unmasked faces. Face responsiveness was defined by comparing the post-stimulus interval (comprising [100,300] ms after stimulus onset) of upright face trials in the unmasked visual task with the pre-stimulus interval (comprising [-200,0] ms relative to stimulus onset) of trials from any category in the same task. The linear separability between these two sets of trials was estimated using RLSC over spectro-temporal representations of the ECoG signals as described above, considering log power at 4x11 (time,frequency) points for each trial, sampled from a uniform grid in the interval [100,300] ms (for the post-stimulus set) or [-200,0] ms (for the pre-stimulus set) relative to stimulus onset x [0,200] Hz, as the input to the classifiers. An electrode was considered to be face-responsive if its decoding accuracy A’ was significant at p<0.01.

#### Comparison between different decoding analyses

In this article, we consider several decoding analyses on neural activity that either contrast an upright face with an inverted face or a non-face stimulus (in both masked and unmasked conditions), or a more visible face with a less visible face (in the masked condition). We hypothesized that the brain loci that are responsible for the generation of conscious experiences of upright faces would exhibit similar levels of discriminability across these different decoding analyses. In order to assess the degree of similarity between different decoding analyses, we computed the Pearson correlation ρ between A’ values for every pair of decoding analyses over face-responsive electrodes, separately for those implanted in the ventral and lateral temporal cortex.

For each pair of decoding analyses and for each brain region, we performed two different statistical tests. First, we tested whether the correlation was significant (against the null hypothesis of nonsignificant correlation) using a permutation-based method: a null distribution of correlation values was constructed by shuffling electrode identity independently for each decoding analysis, calculating the resulting correlation among A’ values and repeating this procedure *N*_perm_=1000 times. An empirical correlation value was considered to be significantly positive (negative) at a significance level p (p=0.05, 0.01, 0.001) if it exceeded (preceded) the 1-p (p) percentile of the corresponding null distribution.

Second, we tested for a significant effect of region label (against the null hypothesis of no effect of region label, i.e., ventral or lateral), again using a permutation-based method. A null distribution of pairwise correlations was constructed by iterating *N*_perm_=1000 times the following procedure: we randomly chose *N*_*x*_ (*x*=ventral, lateral) of (A’_*i*_,A_*j*_) pairs from the pooled set of (A’_*i*_,A_*j*_) pairs (comprising both ventral and lateral electrodes), where *N*_*x*_ is the number of (A’_*i*_,A_*j*_) pairs for region x, and the corresponding Pearson correlation was computed. Then, each correlation coefficient was considered to be higher (lower) than expected by chance (that is, if region labels were irrelevant) at a significance level p (p=0.05, 0.01, 0.001) if it exceeded (preceded) the 1-p (p) percentile of the corresponding null distribution.

In order to visualize the patterns of similarity between different decoding analyses, we performed multidimensional scaling (MDS) on the correlation tables, using D = 1 — ρ as a measure of dissimilarity between pairs of decoding analyses. MDS enables one to represent the original, high-dimensional data (corresponding to one dimension for each decoding analyses considered) in a lower dimensional space (here, two-dimensional) while approximately conserving the relative distances (here, similar patterns of decoding accuracy across electrodes) between data points (Cox and Cox, 2000).

## Results

### Neuronal responses to objective and subjective attributes of visual stimuli

Our electrophysiological data set comprised 1071 bipolar channels (187, 171, 219, 228, 266 from subject 147, 153, 154, 168, 178) from the ventral and lateral temporal cortex of 5 subjects. Out of these, 271 channels were face-responsive (82, 57, 53, 53, 26 from subject 147, 153, 154, 168, 178, see subsection “Face responsiveness” in Methods for the definition of face responsiveness).

A subset of face-responsive channels exhibited spectral power responses that differed between categories in the unmasked task, and between intervals that contained a face image and those that did not in the CFS task. Most of these channels exhibited spectral power responses to CFS face intervals that were modulated by both physical aspects of presented stimuli (i.e. the contrast of a target face), as well as by subjective perception of those same stimuli (i.e. the visibility rating), as in the case of the example ventral electrode shown in Fig. 3.

**Figure 3.**
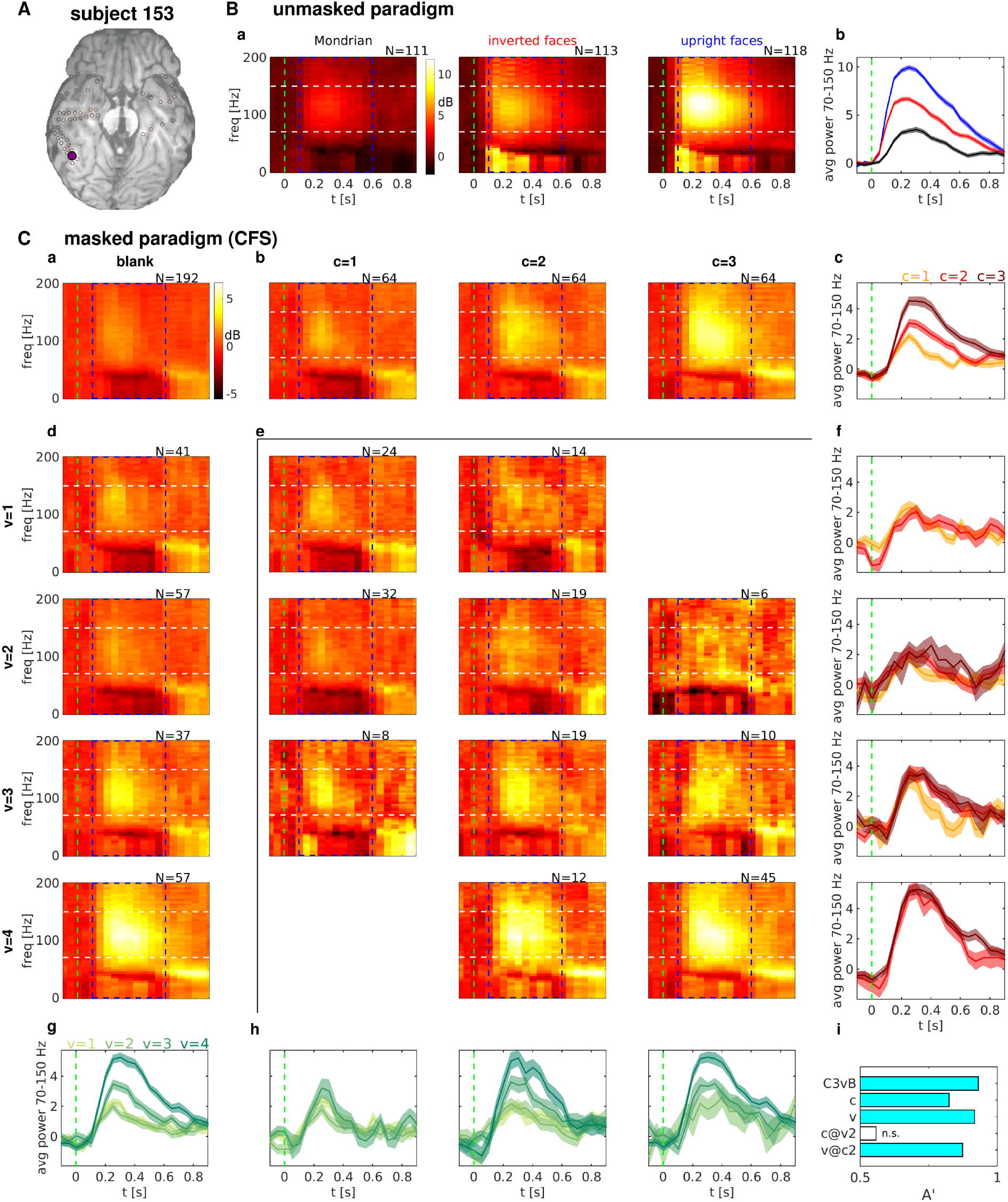
Spectral power responses in the unmasked and CFS experiment for an example electrode. (A) Location of the example electrode. (B) Spectral power responses to some stimulus categories in the unmasked task. (a) Average spectrograms for Mondrian patterns (left), inverted faces (middle) and upright faces (right). (b) Band-limited power (BLP) signals obtained by averaging the corresponding spectrograms in a) over the frequency range delimited by white dashed lines. (C) Spectral power responses in the CFS task. (a) Average spectrogram in “blank” intervals. (b) [d]: Average spectrograms in “face” intervals with different face contrast values [visibility ratings], increasing along columns [rows]. (c) [g]: BLP signals obtained by averaging the corresponding spectrograms in b [d] over the frequency range delimited by white dashed lines. (e) Average spectrograms in “face” intervals corresponding to a given face contrast value (*c*, varying over columns) and visibility rating response (v, varying over rows). Only (*c,v*) pairs resulting in at least 5 trials are shown. (f) [h]: BLP signals obtained by averaging the corresponding spectrograms in e) over the frequency range delimited by white dashed lines. BLP signals with the same visibility rating [face contrast value] but different face contrast value [visibility rating] are shown in the same panel, with visibility [contrast] increasing along rows [columns]. (i) Decoding accuracy A’ for some decoding analyses. C3vB: *c*=3 trials vs. blank trials. *c*: *c*=1 vs. *c*=2,3. v: *v*=1,2 vs. *v*=3,4. *c@v*2: (*c*=1,v=2) vs. (*c*=2,3,v=2). *v@c*2: (*c*=2,*v*=1,2) vs. (*c*=2,*v*=3,4). Cyan bars indicate significant decoding accuracy (p<0.001). Blank bars indicate non-significant decoding accuracy (p>0.05). The number of trials averaged for each condition is indicated on top of each spectrogram. Dash blue rectangles indicate the time-frequency region that has been considered for the decoding analyses. Differences in log power with respect to a pre-stimulus baseline ([-500,0] ms) are shown for the ease of visualization. The green dash vertical lines indicate flash onset. Shaded areas in the BLP signals indicate s.e.m. across trials.

For this electrode, the presentation of an upright face evoked a stronger response than the presentation of an inverted face, which in turn evoked a stronger response than the presentation of a Mondrian pattern in the unmasked task (Fig. 3B). The spectral response exhibited an increase in power located mostly in the high-gamma band (70-150 Hz) and in the alpha-beta band (4-40 Hz). In the CFS task, increasing face contrast values resulted in increases in power mostly in the high gamma band (70-150 Hz) and in the time window from 200 to 500 ms after stimulus onset, with a complex spectral power response in the alpha-beta band characterized by an early attenuation in power decrease followed by a late increase (Fig. 3C;b,c). A very similar pattern of spectral responses could also be observed for increasing visibility ratings (Fig. 3C;d,g), which prompts the question of whether spectral changes due to increasing contrast can be explained, and to which degree, by changes due to increasing visibility rating.

Indeed, averaging spectral responses separately for each pair of contrast value c and visibility rating v shows that spectral changes due to increased contrast at a fixed visibility rating are negligible, while spectral changes due to increased visibility rating at a fixed contrast value can still reliably distinguish different visibility ratings (Fig. 3C;e,f,h,i). This result suggests that, for this electrode, the pattern of spectral changes in response to increasing contrast values can be explained almost completely by changes in visibility ratings, i.e. by changes in subjective perceptual state.

While the electrode shown in Fig. 3 exhibited spectral power responses in both the high-gamma and the alpha-beta bands, other electrodes exhibited different spectral responses. For example, one electrode from the lateral cortex of the same subject exhibited spectral responses to face images of increasing contrast that were almost completely confined to the high-gamma band (Fig. S2), while another electrode from the ventral cortex of another subject showed mostly spectral power increases in the alpha-beta band (Fig. S3).

In the present study, we aimed to assess neuronal responses that are informative of the presented stimuli or the resulting perceptual state regardless of the spectral range in which they are observed. Hence, in the next section, we will present a systematic quantification of the information about stimuli or perceptual states using a multivariate decoding analysis that combines spectral power sampled from a uniform grid in the interval [100,600] ms after stimulus onset x [0,200] Hz, indicated by a blue dash rectangle in the spectrograms of Fig. 3.

### Neuronal correlates of conscious visual awareness

In order to investigate the neuronal correlates of objective physical stimulation, and compare them with the neuronal correlates of subjective phenomenal experience, we performed a set of decoding analyses for each face-responsive electrode. In particular, we evaluated the linear separability, as measured by the performance of a set of binary RLSC classifiers (see Methods for details), between pairs of subsets of CFS intervals. We considered the following contrasts: *i*) face intervals with face contrast level equals three vs. blank intervals (*c*_3_ *vs. blank*); *ii*) high visibility face intervals vs. low visibility face intervals, grouping across all contrast levels (v); and iii) high visibility face intervals vs. low visibility face intervals, considering only trials with a fixed level of face contrast, namely the “threshold face constrast” as defined in section “Behavioral Analysis”, corresponding to intermediate objective performance (*v@c*_2_ or *v@c*_3_ depending on the subject), which we denote as *c*_thr_.

These three contrasts constitute a gradual shift from a criterion defined exclusively by extrinsic factors (*c*_3_ vs. *blank*), to a criterion defined exclusively by the intrinsic, subjective phenomenal experience, in the absence of any change in the physical property of the stimulus (*v@C*_thr_).

The results from these three decoding analyses for an example subject are shown in Fig. 4A. The comparison between a decoding analysis specified by extrinsic factors (*c_3_ vs. blank*) and one specified by subjective perception (*v*) reveals a great degree of overlap between the brain areas that discriminate these two pairs of conditions: the electrodes that differentiate high contrast face intervals from blank intervals are the same ones that differentiate clearly seen face intervals from poorly seen or unseen face intervals. Plotting A’_c3vs_.blank versus A’_*v*_ reveals a strong correlation between the two decoding analyses, with stronger correlation observed among the most discriminant electrodes (Fig. 4B, top panel).

**Figure 4.**
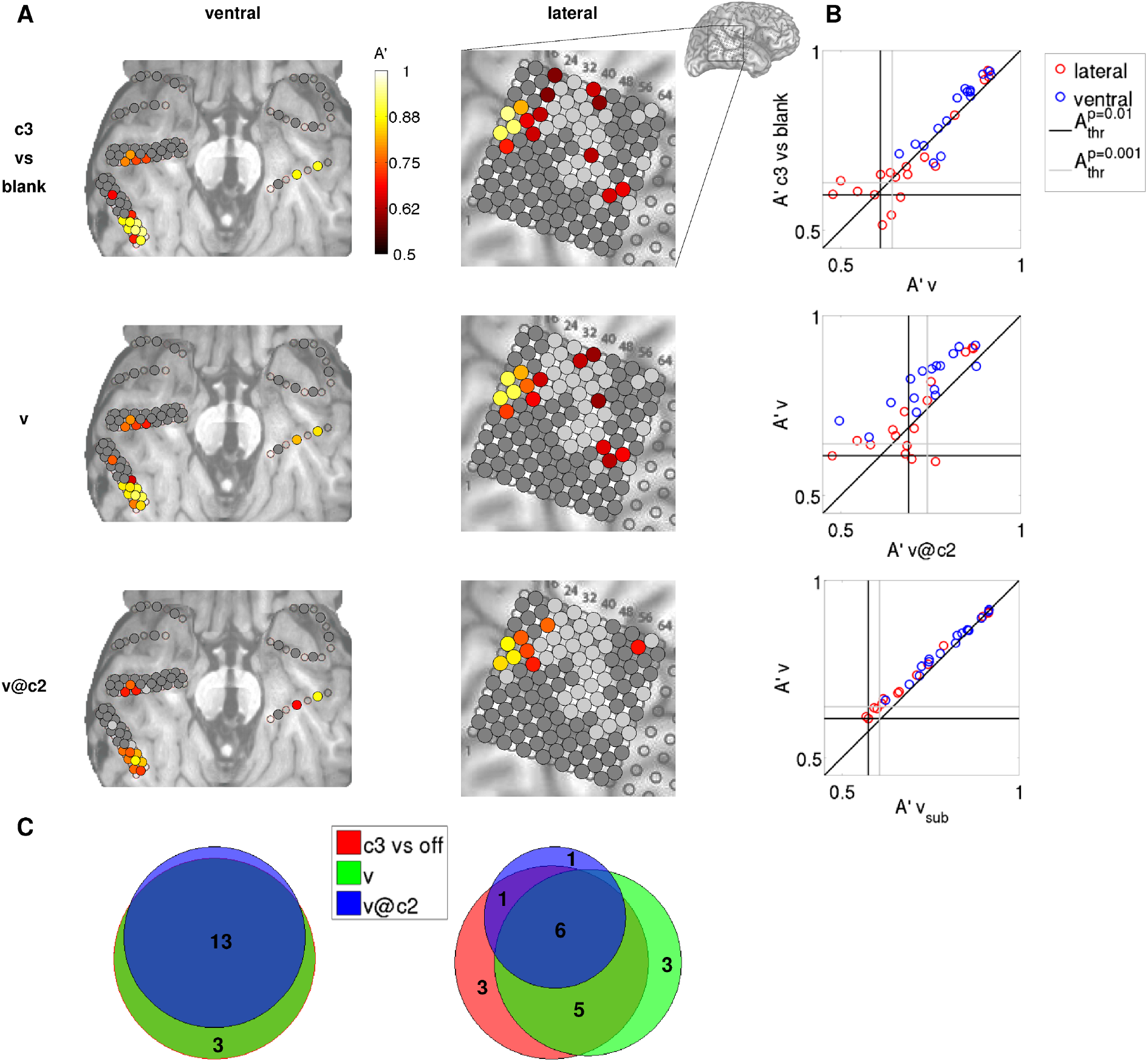
Distilling the neuronal correlates of conscious visual perception: from physical attributes of stimuli to subjective phenomenology. (A) Decoding accuracies A’ for each electrode in the example subject 153 are shown color-coded on the ventral (left) and lateral (right) brain images. The decoding analyses shown progress from a contrast specified by physical stimuli (c3 *vs. blank,* top row), to a contrast defined by subjective phenomenology, but contaminated by external factors (i.e., face contrast, v, middle row), to a purely subjective contrast (*v@c*_2_, bottom row). Only decoding accuracies that are significant at p<0.01 (uncorrected for multiple comparisons) are shown. Dark gray indicates non face-responsive electrodes; light gray indicates electrodes that are face-responsive but do not exhibit significant decoding accuracy. Decoding accuracies A’ for each electrode and for each remaining subject are shown color-coded on the anatomical images in Fig. S4. (B) Relationships between pairs of decoding analyses for each electrode that exhibits a significant A’ in either one of the two decoding analyses. There is a strong correlation between the decoding accuracy for c_3_ vs. *blank* and v, and between v and *v@c*_2_. v_sub_ indicates the decoding accuracy obtained when decoding visibility using the same number of trials for training and test (for each round of cross-validation) as when decoding *v@c*_2_. The vertical (horizontal) black line indicates the p=0.01 significance threshold for the decoding analysis corresponding to the x (y) axis, averaged over electrodes. The vertical (horizontal) gray line indicates the p=0.001 significance threshold for the decoding analysis corresponding to the x (y) axis. Inset shows lateral brain image, with the area enlarged in the main panels indicated with a rectangle. (C) Venn diagrams showing the number of electrodes that are significant (p<0.01) in one or more of the decoding analyses considered, separately for ventral (left) and lateral (right) electrodes. The sets of ventral electrodes that discriminate *c*_3_ vs. *blank* and v (red and green circles) completely overlap for this subject. *c*_3_ vs. *blank,* red; v, green; *v@c*_2_, blue.

The comparison between otherwise unselected high and low visibility trials, even though defined on the basis of a purely perceptual category, will typically include the contribution of different levels of face contrast, since higher face contrast results in higher visibility ratings (Fig. 2B). Hence, a further distillment of the neuronal correlates of conscious face perception can be achieved by contrasting high vs. low visibility trials at threshold contrast *c*_thr_. This analysis reveals a smaller set of electrodes that is almost completely included in the set specified by the v decoding analysis. In particular, the best *v@c*_thr_ discriminant electrodes correspond with those that best discriminate v (Fig. 4B, middle panel). In order to quantify the effect of the different number of trials that enter the v and *v@c*_thr_ analyses, we performed a decoding analysis discriminating visibility using the same number of trials for training and test (for each round of cross-validation) as when decoding *v@c*_2_ (v_sub_ for subsample). The results are very similar as when decoding visibility without subsampling (Fig. 4B, bottom panel), suggesting that the decrease in decoding accuracy generally observed when discriminating *v@c*_t_hr (with respect to discriminating v) is a genuine result of the decreased discriminability between the two classes, and is only minimally affected by the different number of trials. For the subject depicted in Fig. 4, the degree of overlap between the sets of discriminant electrodes specified by these three decoding analyses (c_3_ *vs. blank, v* and *v@c*_2_) is very high for both ventral and lateral electrodes, albeit higher for ventral electrodes (Fig. 4C).

Other subjects exhibited a similar trend (Fig. S4 and 5A), although fewer electrode sites were *v@c*_thr_ discriminant (see also Discussion and Fig. S7). Importantly, the strong correlation between decoding accuracies among the best discriminant electrodes in the *c*_3_ vs. *blank* and v decoding analyses, and in the v and *v@c*_thr_ decoding analyses, was conserved when electrodes were pooled across subjects (Fig. 5A). The substantial degree of overlap between the three sets of discriminant electrodes, with higher overlap for ventral electrodes, was also conserved (Fig. 5B). We will quantify the degree of similarity between different decoding analyses further below (“Comparison between different decoding analyses”).

**Figure 5.**
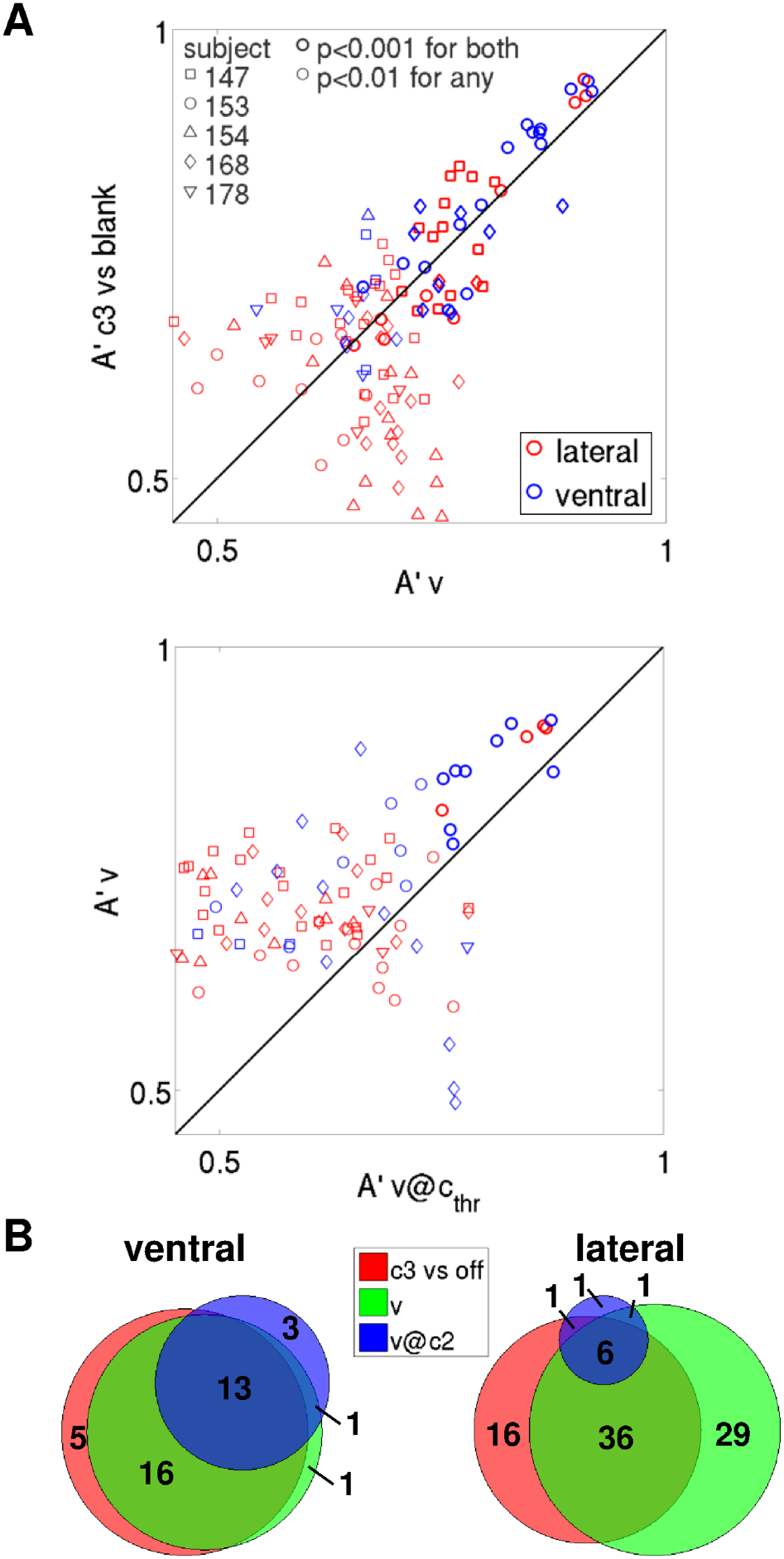
Distilling the neuronal correlates of conscious visual perception: from physical attributes of stimuli to subjective phenomenology. (A) As in Fig. 4B, pooling electrodes across subjects. Decoding accuracies A’ for each electrode and for each remaining subject are shown color-coded on the anatomical images in Fig. S4. (B) As in Fig. 4C, pooling electrodes across subjects.

To further characterize the modulation of neuronal responses attributable to physical attributes of stimuli (i.e., face contrast), and compare it with the modulation attributable to subjective visibility, we also performed a decoding analysis that contrasted low contrast trials vs. high contrast trials (c decoding), after grouping adjacent contrast levels using a median split of the data.

All subjects, and in both ventral and lateral cortices, presented several electrodes that reliably differentiated low vs. high contrast trials (*c* decoding, shown in Fig. 6A for the example subject 153, left half of the bisected disks), as well as low vs. high visibility trials (v decoding, Fig. 6A, right half of the bisected disks). Decoding accuracy is generally higher for decoding visibility than contrast. In the case of the example subject 153, the cumulative probability density of decoding accuracy among face-responsive electrodes is consistently lower for *v* than for *c* in both ventral and lateral electrodes (p<3.2 · 10^-4^ in both cases, Kolmogorov-Smirnov test), which corresponds to a probability density shifted to the right (that is, towards higher decoding accuracies) when decoding v in comparison with decoding *c* (Fig. 6C, left panel). More importantly, the highest decoding accuracy is obtained when decoding visibility, rather than contrast (max(A’_*v*_)=0.917 among ventral electrodes, max(A’_*v*_)=0.911 among lateral electrodes; max(A’_*c*_)=0.825 among ventral electrodes, max(A’_*c*_)=0.82 among lateral electrodes). Also, 93.75% (19.51%) of face-responsive ventral (lateral) electrodes have a *v* decoding accuracy that is significant at p<10^-5^, as compared to 56.25% (9.76%) of face-responsive ventral (lateral) electrodes that have a *c* decoding accuracy with the same level of significance (Fig. 6C, left panel; note that similar results hold if considering less conservative significance thresholds). For this subject, decoding accuracy among ventral electrodes is higher than among lateral electrodes for both c and v decoding (p<1.9 · 10^-13^ in both cases, Kolmogorov-Smirnov test), although the highest accuracy observed in each region is similar.

**Figure 6.**
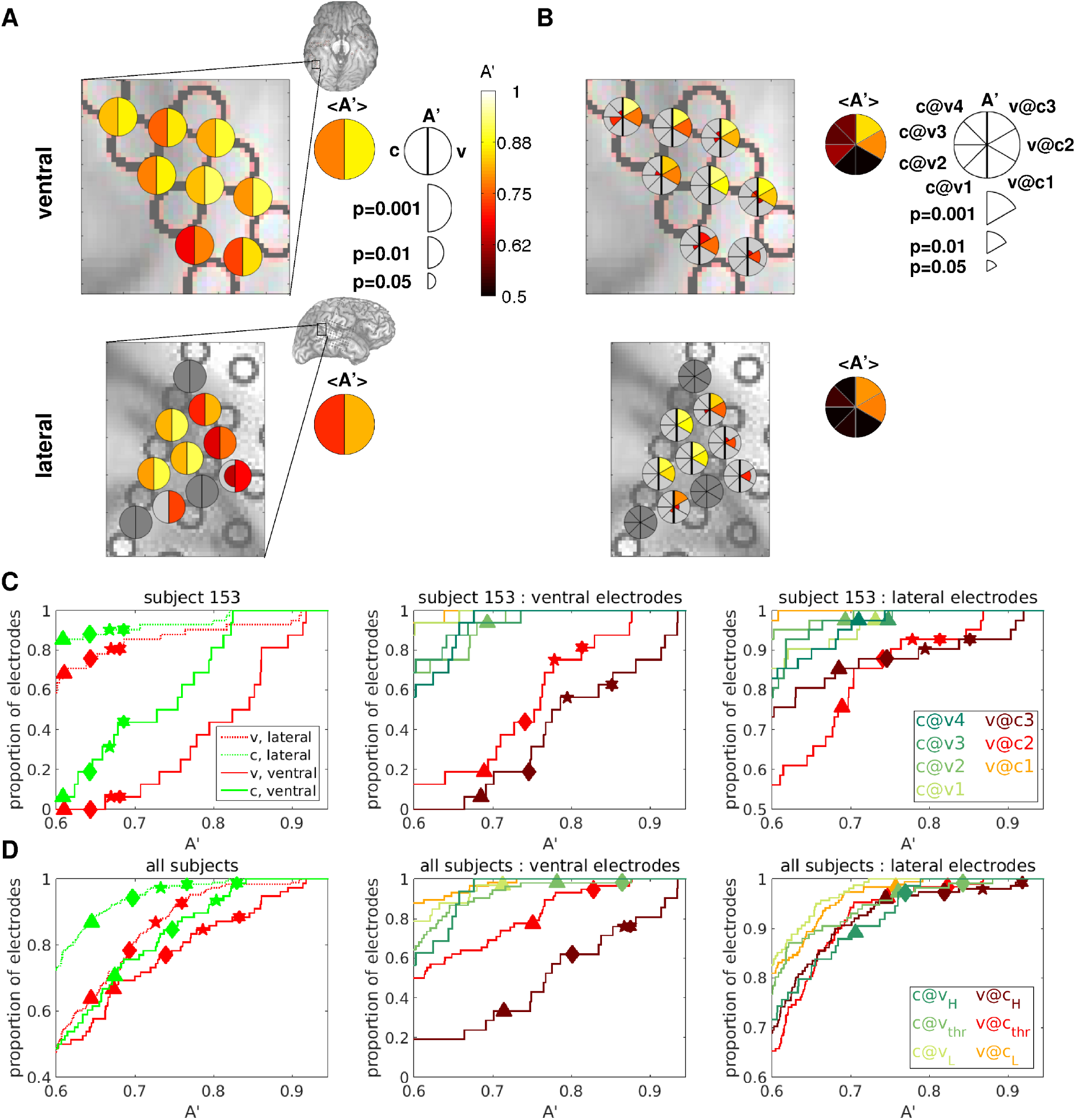
Subjective visibility can be decoded better than physical contrast. (A) Decoding accuracies A’ for a selected group of electrodes from an example subject are shown color-coded on the ventral (top) and lateral (bottom) brain images. A’_c_ are indicated in the left half of the bisected disks, A’_v_ in the right half. Only decoding accuracies that are significant at p<0.05 (uncorrected for multiple comparisons) are shown, with larger symbol size indicating higher significance. Brain images show the areas that are enlarged in the main panels. Summary disks indicate the average A’ among the face-responsive electrodes in the selected area. (B) As in A, but with A’*_c_@_v_* indicated in the left sectors of the pie charts, A’*_v_@_c_* in the right sectors, as shown in the legend. (C), (D): Cumulative probability density functions of decoding accuracy over the populations of ventral and lateral electrodes for an example subject (C), and pooling over all subjects (D). Only results from decoding analyses with at least 10 trials in the least populated class are shown. Symbols indicate threshold A’ at several significance levels for each classification considered, obtained by permutation. Symbols are only shown if there is at least one electrode that is significant for the corresponding decoding analysis and significance level. Triangles: p=0.01; Diamonds: p=0.001; Stars: p=0.0001; Hexagons: p=0.00001. Single-subject cumulative probability density functions of decoding accuracy for the remaining subjects are shown in Fig. S5.

The cumulative probability density functions of decoding accuracies for the remaining subjects are also consistent with the hypothesis that decoding visibility results in higher accuracy than decoding contrast (Fig. S5). In particular, the cumulative probability density of decoding accuracies among ventral electrodes is significantly lower for v than for c in 3/5 subjects (p<0.007, Kolmogorov-Smirnov test), yielding non-significant results in subject 168, while the relationship is inverted in subject 178 (i.e., the cumulative probability density of decoding accuracies is significantly lower for c than for v, p=2.8 · 10^-4^, Kolmogorov-Smirnov test). Note, however, that this subject only presents one ventral electrode that is marginally significant when decoding c (p<0.01), hence the cumulative probability densities of decoding accuracies among ventral electrodes for both c and v are composed mostly of non-significant A’ values for this subject (Fig. S5). Among lateral electrodes, the cumulative probability density of decoding accuracies is significantly lower for v than for c in 3/5 subjects (p<8.7 · 10^-4^, Kolmogorov-Smirnov test), yielding marginally significant results in the other two subjects (154 and 168, p=0.019 and p=0.028, respectively).

Importantly, similar results are obtained when pooling electrodes across subjects: the cumulative probability density (cpdf) of decoding accuracy among face-responsive electrodes is consistently lower for v than for c in both brain areas, and it is consistently lower in ventral than lateral electrodes for both decoding analyses (Fig. 6D, left panel). These relationships reach statistical significance in the case of the v cpdf being lower than the c cpdf among lateral electrodes (p=4 · 10^-12^, Kolmogorov-Smirnov test), and in the case of cpdfs being lower among ventral than among lateral electrodes for both c and v decoding (p=1.4 · 10^-8^ for the former, p=0.0068 for the latter, Kolmogorov-Smirnov test). The v cpdf is consistently lower than the c cpdf among ventral electrodes as well, but the relationship does not reach statistical significance (p=0.12, Kolmogorov-Smirnov test). Note, however, that the maximum difference between the v and the c cpdfs is observed for low, non-significant values of the decoding accuracy in the set of lateral electrodes, while it is observed for high and strongly significant values of the decoding accuracy in the set of ventral electrodes (for A’∽0.83, Fig. 6D, left panel). Also, 11.54% (7.25%) of face-responsive ventral (lateral) electrodes have a v decoding accuracy that is significant at p<10^-5^, as compared to 1.28% (1.55%) of face-responsive ventral (lateral) electrodes that have a c decoding accuracy with the same level of significance (Fig. 6D, left panel).

To further dissociate the modulation of neuronal responses attributable to face contrast and subjective visibility separately, we performed a set of decoding analyses for each electrode. In particular, we calculated the linear separability A’ between low and high contrast trials corresponding to a fixed value of subjective visibility (*c@v* decoding, for each of the four visibility ratings), and the linear separability A’ between low and high visibility trials corresponding to a fixed face contrast value (*v@c* decoding, for each of the first three contrast values). In all cases, we performed binary classifications after grouping adjacent contrast values (for *c@v* decoding analyses) or visibility ratings (for *v@c* decoding analyses).

The results of these analyses for a set of visibility and contrast discriminant electrodes from an example subject are shown in Fig. 6B. When considering a fixed value of subjective visibility, face contrast was no longer decodable, and decoding accuracies dropped to non-significant or marginally significant levels (*c@v* decoding, left sectors). Conversely, when considering a fixed value of face contrast high enough to enable above chance performance (*c*=3 or *c*=2), subjective visibility was still decodable with very high accuracy (*v@c*_3_ and *v@c*_2_, first two right sectors from the top).

In the example subject, the remarkably stronger modulation of neuronal activity due to visibility at fixed contrast, rather than due to contrast at fixed visibility, is also reflected at the electrode population level in the cumulative probability density functions (cpdfs) of decoding accuracies for ventral and lateral electrodes shown in Fig. 6C (middle and right panels). While several electrodes in both ventral and lateral areas display decoding accuracies that are significant at p<10^-5^ for both *v@c*_2_ and *v@c*_3_, *v@c*i and *c@v* decoding analyses only result in at most one electrode that is significant at 10^-3^<p<10^-2^, as expected by chance. This trend is also observed when pooling electrodes across the population of subjects (Fig. 6D, middle and right panels). However, the accuracy of *c@v*_thr_ and *c@v*_H_ decoding is also high, especially among lateral electrodes, mostly driven by the contribution of *c@v*_4_ from subjects 154, 168 and 178 (Fig. S5, see also Discussion).

It is worth noting that subjects 154, 168 and 178 were overconfident in their visibility ratings. In particular, their performance when reporting the highest visibility rating *v*=4 was not greater than 91% (91%, 87% and 72%, for subject 154, 168 and 178), as opposed to 100% for subjects 147 and 153 (Fig. 2D). Hence, it is possible that they responded with the highest visibility rating even if their perception of the face was not completely clear. If this were the case, the high accuracy in *c@v*_4_ decoding they exhibit might reflect different degrees of subjective face perception, which is expected to covary with face contrast in v=4 trials in overconfident subjects. In accordance with this interpretation, most electrodes with significant accuracy in *c@v*_4_ decoding also display significant *v* or *v@c*_thr_ decoding accuracy.

### Distilling the neuronal correlates of conscious visual awareness by combining masked and unmasked presentation of similar images

The comparison between seen and unseen trials, even in conditions of equal physical stimulation, is not guaranteed to reveal the core neuronal correlates of a specific conscious perception. As pointed out previously (Aru et al., 2012b; de Graaf et al., 2012; Miller, 2007; van Boxtel and Tsuchiya, 2015; Tsuchiya et al., 2015), such comparison is also expected to expose neuronal activities that can facilitate or hinder the perceptual experience of a faint stimulus, as well as neuronal activities that are related to the motor act of perceptual report.

Hence, in addition to the masked paradigm, we also considered an unmasked paradigm, where images of faces and other categories were shown at the fixation point for 500 ms without presenting any competing stimuli or any perceptual mask, guaranteeing an effortless and vivid perception of the presented objects. As opposed to the masked paradigm, where identical stimuli could elicit different perceptual outcomes, the unmasked presentation is expected to result in a one-to-one correspondence between physical stimuli and perceptual states.

In order to identify the brain loci that discriminate between images of upright faces and other categories, we performed two decoding analyses for each electrode: i) a more generic analysis, contrasting upright face trials with trials where other categories were presented (5 subjects), and ii) a more specific analysis, contrasting upright face trials with inverted face trials (3 subjects).

The decoding accuracies resulting from these analyses, for the same electrodes shown in Fig. 6A, are color-coded in the two left quarters of the pie charts in Fig. 7A, with the more generic analysis (upright face vs. other categories) shown on top, and the more specific analysis (upright face vs. inverted face) shown on bottom. The two right quarters show the decoding results from the masked paradigm for comparison, with a more generic analysis (v) shown on top, and a more specific analysis (*v@c*_2_) shown on bottom.

**Figure 7.**
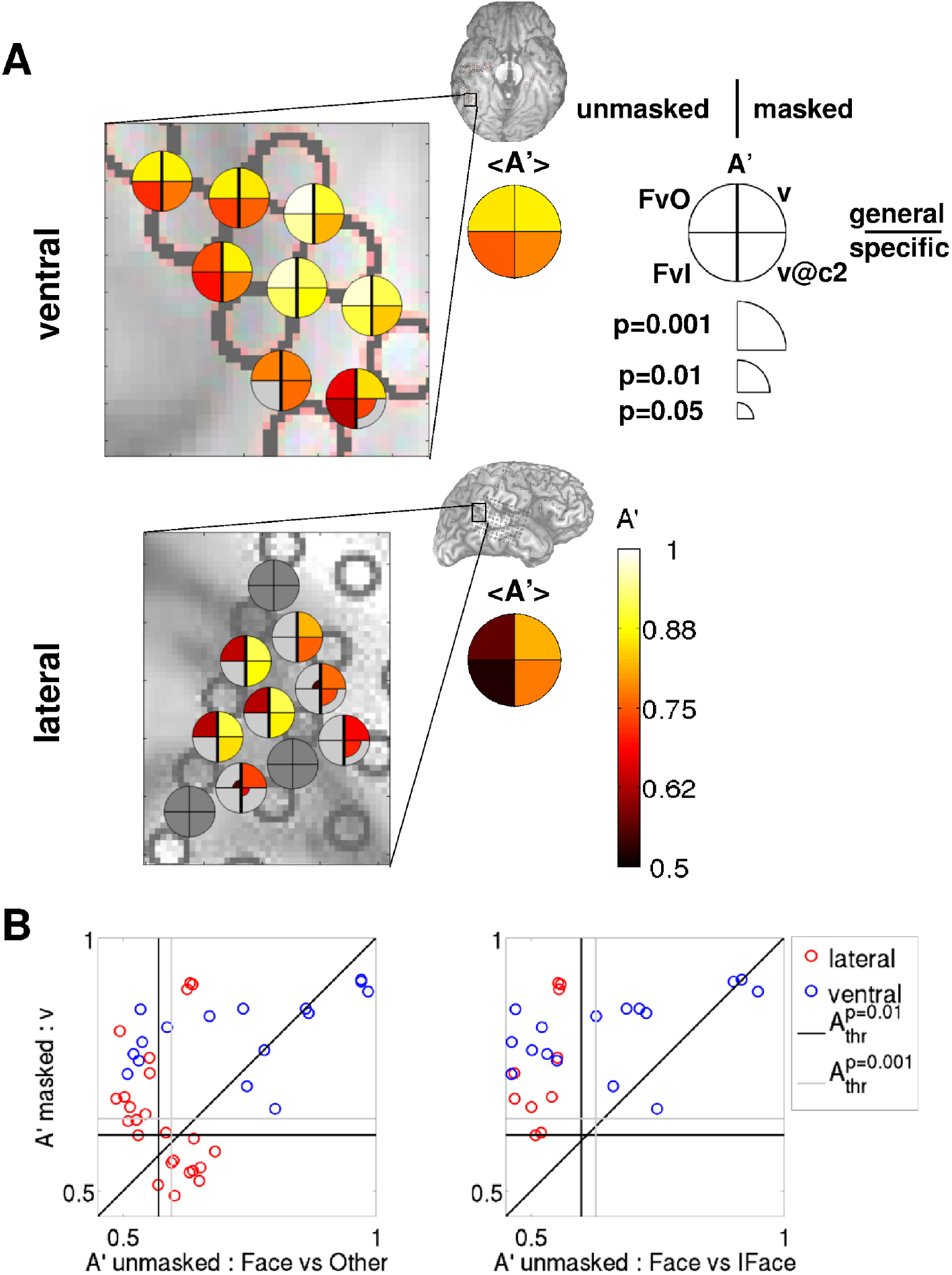
Comparison between decoding visibility in the masked paradigm and decoding image category in the unmasked paradigm: both ventral and lateral areas can discriminate visibility, but only ventral areas can discriminate upright from inverted faces. (A) Decoding accuracies A' for a selected group of electrodes from the example subject 153 (the same as shown in Fig. 6A,B) are shown color-coded on the ventral (top) and lateral (bottom) brain images. Accuracies in upright face decoding using the unmasked protocol are shown in the left quadrants (top: upright face vs. other categories; bottom: upright vs. inverted face), accuracies in visibility decoding using the masked protocol are shown in right quadrants (top: v; bottom: *v@c*_2_), as indicated in the legend. Only decoding accuracies that are significant at p<0.05 (uncorrected for multiple comparisons) are shown, with larger symbol size indicating higher significance. Brain images show the areas that are enlarged in the main panels. Summary disks indicate the average A' among the face-responsive electrodes in the selected area. (B) v decoding accuracies in the masked paradigm are plotted against the corresponding decoding accuracies when discriminating upright faces in the unmasked paradigm for each electrode from the example subject that is significant at p<0.01 in at least one of the decoding analyses for each pair of decoding analyses. A more generic unmasked decoding analysis is shown in the left panel (upright faces vs. other categories), a more specific in the right panel (upright vs. inverted faces). The vertical (horizontal) black [gray] line indicates the p=0.01 [0.001] significance threshold for the decoding analysis corresponding to the x (y) axis, averaged over electrodes.

While both ventral and lateral regions can discriminate upright faces versus other categories (top-left quadrant), with higher accuracy in ventral than in lateral regions, only ventral loci can discriminate between upright and inverted faces (bottom-left quadrant).

This remarkable difference is also evident at the electrode population level when comparing decoding accuracies in the masked paradigm (*v* and *v@c*_thr_) with those in the unmasked paradigm (upright face vs. other categories and upright vs. inverted face) (Fig. 7B and Fig. 8). While both ventral and lateral electrodes can discriminate visibility (*v*) in the masked paradigm (even when the decoding analysis is restricted to the threshold contrast (*v@c*_thr_), see Fig. 4, 5 and 6), as well as upright face vs. other categories in the unmasked paradigm (with greater accuracy in ventral than lateral regions), only ventral electrodes can discriminate upright vs. inverted faces.

**Figure 8.**
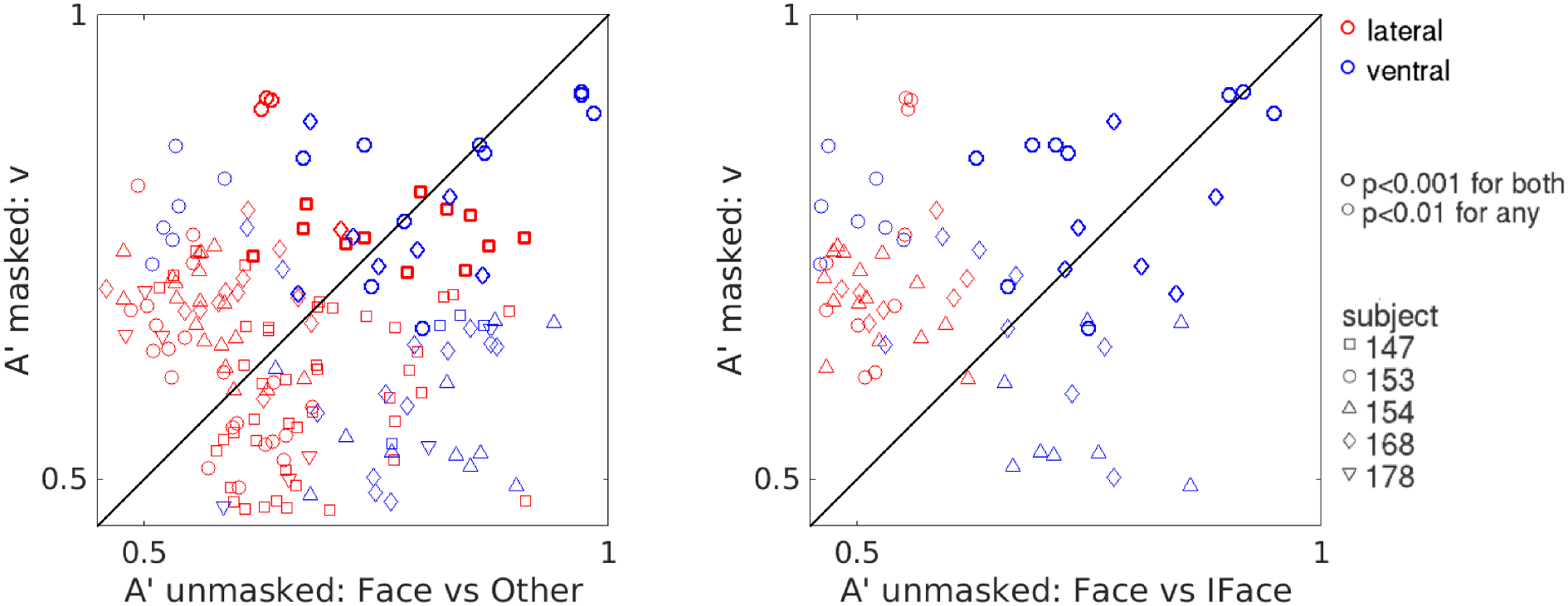
Comparison between decoding visibility in the masked paradigm and decoding image category in the unmasked paradigm across all subjects. As in Fig. 7B, pooling electrodes across subjects. While both ventral and lateral regions can discriminate visibility in the masked paradigm and upright faces from other categories in the unmasked paradigm, only ventral areas can discriminate upright from inverted faces.

In all three subjects for which the upright vs. inverted face decoding analysis is possible, several ventral electrodes display high and very significant (p<0.001) decoding accuracy, and only one lateral electrode shows marginally significant decoding accuracy (0.001<p<0.01), as would be expected by chance considering the number of electrodes that comprise our data set. Ventral electrodes that reliably discriminate upright vs. inverted faces are observed in both right (subjects 153 and 154) and left (subject 168) fusiform gyri.

As shown in the average spectrograms and band-limited power signals in Fig. 3 and Fig. S2, upright faces often elicit the strongest response in both ventral and lateral electrodes. However, the great variability across trials in lateral electrodes prevents accurate discrimination between different categories, and critically, between upright and inverted faces, on a trial-by-trial basis. Conversely, ventral electrodes record stronger activations that are more reliable across trials of any given category, hence enabling accurate discrimination of upright faces.

When searching for the neuronal correlates of conscious perception, we assume that areas belonging to the NCC-core will exhibit activity that discriminates conditions corresponding to different perceptual experiences. While both upright and inverted faces are recognized as “faces”, the corresponding phenomenology is quite different (Rossion and Gauthier, 2002; Maurer et al., 2002). Thus, we expect brain areas corresponding to the NCC-core of upright faces to display neuronal activity that discriminate these two classes of stimuli accordingly. The high discriminability for high versus low visibility trials observed in some ventral electrodes (which is preserved even when only trials at threshold contrast are considered), and most prominently in those located in the fusiform gyrus, together with the high discriminability between upright versus inverted faces observed in those same electrodes, suggest that the corresponding brain loci are likely to be part of the core network that generates conscious experiences of upright faces.

### Comparison between different decoding analyses

All the decoding analyses we considered in this work (with the exception of *c@v*) contrast a condition of conscious upright face perception versus a condition of less conscious upright face perception or non upright face perception. Thus, it is reasonable to hypothesize that the brain loci that are responsible for the generation of conscious experiences of upright faces would exhibit similar levels of discriminability across these different decoding analyses. Hence, we systematically compared 7 different decoding analyses (5 from the masked paradigm, 2 from the unmasked paradigm) by calculating the Pearson correlation ρ among every pair of decoding accuracies, for the set of ventral and lateral face-responsive electrodes separately (Fig. 9A).

**Figure 9.**
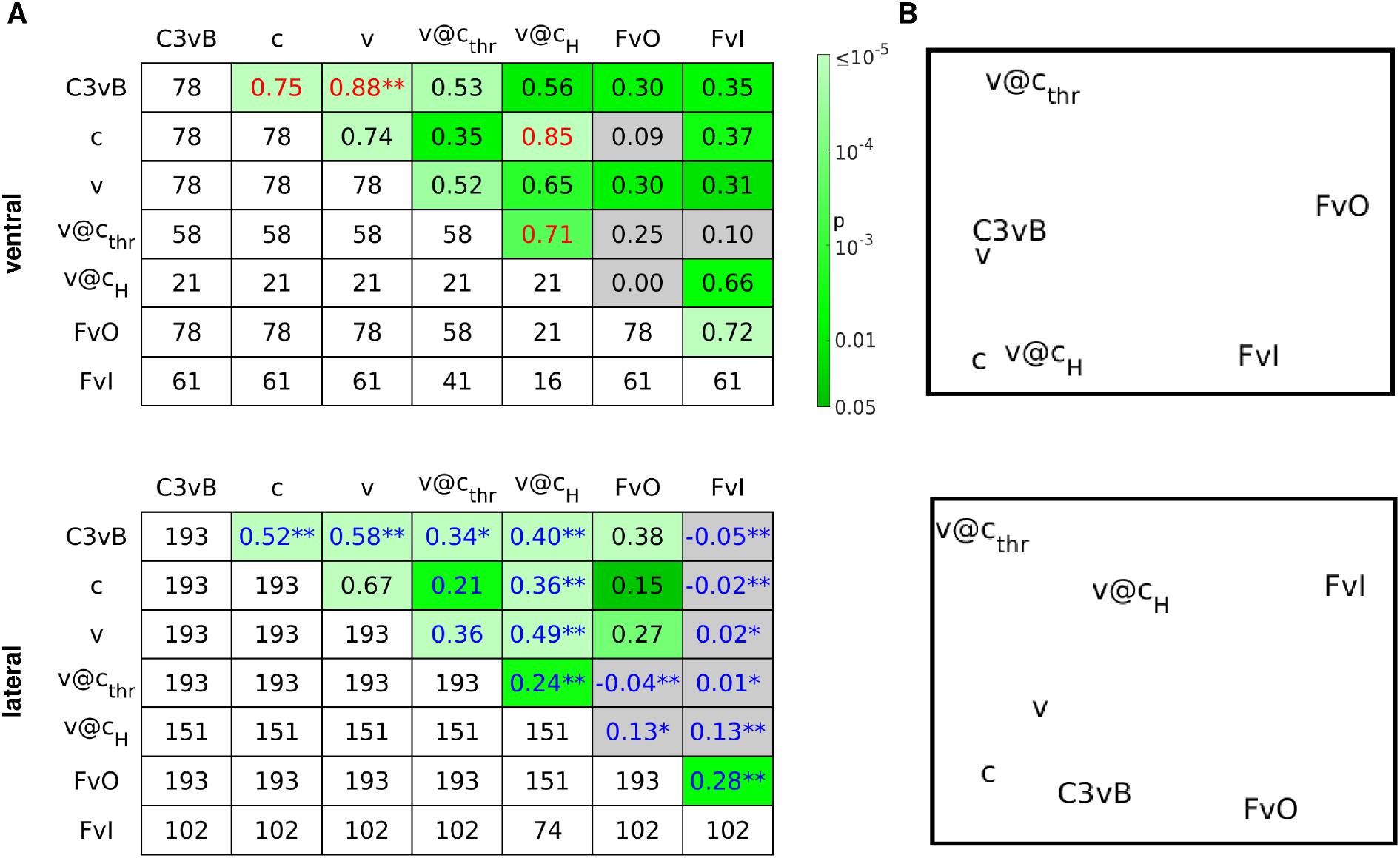
Similarity structure across different decoding analyses. Correlation analysis between A’ values obtained in different decoding analyses for ventral (top) or lateral (bottom) electrodes. (A) Pearson correlation coefficients for each pair of decoding analyses are shown in the upper right part of the table, p-values for the null hypothesis of no correlation are shown color-coded from dark green (p<0.05) to pale green (p<10^-5^). Gray corresponds to p>0.05. Correlation coefficients are printed in red (blue) if higher (lower) than what would be expected by chance if region labels were irrelevant, at p<0.05 (^*^: p<0.01, ^**^: p<0.001, uncorrected for multiple comparisons). Entries in the lower left part of the table show the number of electrodes that enter the corresponding correlation analysis. Diagonal entries indicate the number of face-responsive electrodes for which the assessment of the corresponding decoding accuracy A’ was possible. Correlation analyses were performed separately for ventral (top) and lateral (bottom) electrodes. (B) MDS representations of the correlation tables shown in A. C3vB: *c_3_ vs. blank* ; FvO: upright face vs. other categories; FvI: upright vs. inverted face.

Consistently with the analyses shown in Fig. 4 and 5, we observed high and significant correlation between *c_3_ vs. blank* and *v* decoding (0.88 and 0.58 among ventral and lateral electrodes, respectively) as well as between v and *v@c*_thr_ (0.52 and 0.36) in both ventral and lateral regions. As expected from the results shown in Fig. 7 and 8, we found rather low but significant correlation between decoding v in the masked paradigm and decoding image category in the unmasked paradigm in ventral regions (upright face vs. other categories *FvO* or upright vs. inverted face *FvI*: 0.30 and 0.31). However, the corresponding correlation among lateral electrodes was significant only for FvO (0.27), while for FvI it was equal to 0.02 and not significantly different from zero. In general, correlation coefficients between decoding analyses corresponding to the same paradigm tended to be higher than between decoding analyses corresponding to different paradigms (within paradigm correlations range from 0.35 to 0.88 in ventral regions, and from 0.21 to 0.67 in lateral regions; across paradigm correlations range from 0 to 0.66 in ventral regions, and from −0.05 to 0.38 in lateral regions).

To assess if correlation coefficients were significantly affected by anatomical location (i.e., ventral versus lateral), we performed a set of permutation-based significance tests (see Methods for details). This second-order analysis shows that pairs of decoding analyses applied to the population of ventral electrodes often yield correlation coefficients that are significantly higher than what would be expected if the brain region was irrelevant (4 among 21 pairs tested, indicated with red text in Fig. 9A, top panel).

For example, the accuracy for decoding *c* (a discrimination specified in terms of a physical property of stimuli) is highly correlated with the accuracy for decoding *v@c*_H_ (a purely subjective discrimination) among the set of ventral electrodes (ρ=0.85, significant at p<10^-5^). Hence, ventral electrodes tend to exhibit similar decoding accuracy for the two discriminations and, in particular, ventral electrodes that discriminate c well also tend to discriminate well *v@c*_H_, and vice versa. The same analysis applied to the set of lateral electrode yields a much lower, albeit highly significant, correlation (ρ=0.36, significant at p<10^-5^). Testing for a significant effect of region label reveals that the correlation among lateral electrodes is significantly lower than what would be expected by chance (i.e., if region labels were irrelevant, p<0.001, permutation test). Conversely, the correlation among ventral electrodes is significantly higher than chance (p<0.05), suggesting that decoding accuracies for c and *v@c*_H_ are more similar among the set of ventral electrodes. In addition, a significant positive effect of region label at p<0.001 has been observed among ventral electrodes in the comparison between *c_3_ vs. blank* and *v,* and at p<0.05 between *C*_3_ *vs. blank* and *c*.

Conversely, pairs of decoding analyses applied to the population of lateral electrodes generally yield correlation coefficients that are significantly lower than what would be expected by chance (17 among 21 pairs tested, indicated with blue text in Fig. 9A, bottom panel). In particular, a significant negative effect of region label at p<0.001 has been observed in the comparison between *c_3_ vs. blank* and c, between *c_3_ vs. blank* and v, between *c_3_ vs. blank* and *v@c*_H_, and, critically, in almost every pair comprising *v@c*_H_ or upright vs. inverted face *(FvI*). Hence, electrodes in ventral regions tend to show similar decoding accuracy between two different decoding analyses, while those in lateral regions tend to show more variable decoding accuracy depending on the specific type of decoding analysis performed.

This pattern of results could originate from a large population of lateral electrodes, out of which only a few are part of the core NCC network for face perception and most contribute with non-significant decoding accuracies, and a smaller population of ventral electrodes, out of which many or most are part of the core NCC network. In fact, similar second-order analyses performed on the set of electrodes that are significant in both decoding analyses at different levels of significance yielded correlation coefficients that still tend to be higher among ventral electrodes for most pairs of decoding analyses, but significant effects of region labels are sparser and not always consistent among regions. Future studies with larger samples will be needed to rule out this possibility with greater confidence. It is notable, however, that pairs of decoding analyses including a purely subjective analysis in the masked paradigm (*v@c*_t_hr and *v@c*_H_) and a face discriminant analysis in the unmasked paradigm (upright face vs. other categories *FvO* or upright vs. inverted face *FvI*) still yield much higher correlation coefficients among ventral electrodes, which sometimes reach high levels of significance in the effect of region label (e.g., *v@c*_thr_ and *FvO,* and *v@c*_H_ and *FvO,* both yield significant correlation among ventral electrodes with a positive effect of region label which is significant at p<0.001 if only electrodes that are significant at p<0.01 in both analyses for each pair are considered). This suggests that ventral electrodes actually discriminate different, but similarly related, pairs of conditions in a more similar manner than lateral electrodes.

In order to provide an intuitive representation of the patterns of similarity between different decoding analyses, we performed a multidimensional scaling (MDS) analysis on the correlation tables, considering *D* = 1 — *ρ* as a measure of dissimilarity between pairs of decoding analyses (Fig. 9B). This analysis shows that *c_3_ vs. blank, c* and *v* analyses are very similar to each other in both brain regions, with *c*_3_ *vs. blank* and v particularly close for ventral electrodes. The two analyses of visibility at fixed contrast, *v@c*_thr_ and *v@c*_H_, are also located in the same region of the MDS space, with *v@c*_H_ especially close to c among ventral electrodes. The two decoding analyses from the unmasked paradigm, upright face vs. other categories and upright vs. inverted face, are also similar to each other, especially among the set of ventral electrodes. The upright vs. inverted face decoding results are especially close to *v@c*_H_ among ventral electrodes.

## Discussion

In this work, we presented a progressive distillment of the neuronal correlates of conscious face perception using ECoG recordings in humans during the presentation of both masked and unmasked images of faces and other categories. In the first part of the article, we focused on the masked paradigm and presented a series of decoding analyses that progressed from the identification of the brain loci that discriminate the presence versus the absence of face images (*c*_3_ vs. *blank* decoding), to those that discriminate clearly seen vs. poorly seen or unseen faces regardless of face contrast (v decoding), to those that discriminate clearly seen vs. poorly seen or unseen faces in conditions of equal face contrast (*v@c*_thr_ and *v@c*_H_). In particular, we identified brain areas in both ventral and lateral cortices that reliably differentiate seen versus unseen faces even in conditions of equal face contrast. We observed that subjective visibility is better decodable than physical contrast. Importantly, visibility is still decodable even when only trials at a fixed value of face contrast are considered. On the other hand, contrast values are in general not decodable when considering only trials with a given visibility rating.

In the second part of the article, we also considered a different, albeit related, experimental protocol, where images of faces and other categories were presented unmasked and could be perceived effortlessly. The comparison between the two protocols revealed a critical difference between ventral and lateral brain loci: whereas ventral electrodes located in the fusiform gyrus (FG) could reliably discriminate upright versus inverted faces, electrodes located in the superior temporal sulcus (STS) could only discriminate upright faces versus other categories, but lacked the information required to discriminate upright versus inverted faces.

To relate the different measures of neuronal discriminability in both masked and unmasked procols, we performed a correlation-based similarity analysis across 7 different decoding analyses (5 based on the masked paradigm, 2 based on the unmasked paradigm), each of which constitutes a contrast between an upright face or more conscious face perception versus a non upright face or less conscious face perception. This second-order analysis revealed greater similarity between different decoding analyses in ventral than in lateral regions. Since brain loci that are more closely related with the generation of conscious visual experiences of upright faces are expected to exhibit similar levels of discriminability across these different decoding analyses, this result is also consistent with ventral areas being more closely involved in conscious experiences of upright faces.

### Relationship with previous studies

Taken together, our results suggest a prominent role for ventral areas of the brain, and in particular for the fusiform gyrus, in the generation of conscious face percepts. Our results are consistent with a broad body of literature that assessed the neuronal correlates of face perception using several methodologies, in both clinical and healthy populations.

Prosopagnosia, a neurological condition where the recognition of faces (and other stimuli that involve configural internal templates) is impaired, is associated with lesions in the fusiform gyri (Damasio et al., 1982). While most clinical cases are associated with bilateral damage, cases of prosopagnosia associated with lesions confined to the right hemisphere have also been reported, suggesting a prominent role of the right fusiform gyrus for normal face processing (Landis et al., 1986; De Renzi et al., 1994). However, more recent studies demonstrated the importance of a broader network for normal face perception (Atkinson and Adolphs, 2011), with prosopagnosia sometimes occurring in patients with intact right FG but lesions in the right inferior occipital cortex (Rossion et al., 2003; Schiltz et al., 2006).

A broad wealth of studies, mostly based on fMRI, support a dissociation between the STS and the FG, with the STS being involved mostly in processing changeable aspects of faces, such as emotional expressions, and the FG with more stable attributes of faces, such as identity (see, for example, (Andrews and Ewbank, 2004; Kanwisher and Yovel, 2006)). However, the results from several recent studies challenge this traditional view. It has been shown that information on emotional expression is also present in the FG, and it is actually better decodable from ECoG electrodes located in this area rather than in the STS (Tsuchiya et al., 2008). Also, some studies reported that face identity is better (Kriegeskorte et al., 2007) or equally well decodable (Nestor et al., 2011; Anzellotti et al., 2014) from anterior regions of the inferotemporal cortex, rather than from the FG (but see (Axelrod and Yovel, 2015) for a study that found higher decoding accuracy for famous face identity in the FG than in the anterior inferotem-poral cortex). Furthermore, FG activity is also affected by several lower-level stimulus features, such as contrast, size, orientation and position (Yue et al., 2011; Xu et al., 2009), while neuronal activity in the anterior inferotemporal cortex is mostly invariant to identity-irrelevant properties (Anzellotti et al., 2014; Anzellotti and Caramazza, 2015).

It has been argued that any difference between neuronal activity in STS and FG, as measured by ECoG electrodes, is necessarily confounded by different signal-to-noise ratios (SNRs) in the two areas, since cortical surface electrodes are less well suited to recording activity from within a sulcus (e.g. the STS) than from the surface of a gyrus (e.g. the FG) (Said et al., 2011). While we believe that differences in SNR between these two regions are possible and might have affected our results (in particular, the upright vs. inverted face decoding results in the unmasked paradigm, Fig 7 and 8), we believe it is unlikely that they can completely explain them, since some electrodes in both STS and FG exhibit similar SNR when decoding visibility in the masked paradigm (Fig. 3, S2 and 4). However, we believe that this is an important caveat of the current study and one that deserves to be investigated in detail, preferably through simultaneous ECoG and microelectrode recordings.

Many studies aimed to identify the brain regions whose activity most closely matches the behavioural effects of specific face manipulation. Among these, the face inversion effect (FIE) is one of those that have been most thoroughly studied. This phenomenon consists in the perceptual impairment in response to inverted vs. upright faces that is disproportionally large if compared with the inversion of other visual stimuli (reviewed in (Rossion and Gauthier, 2002; Maurer et al., 2002)), and is widely considered as evidence for the holistic or configural (as opposed to featural or part-based) perception of faces.

An fMRI study reported that the BOLD activity in the FG, but not in the STS, correlated with the behavioral FIE across subjects, and showed greater sensitivity to face identity when faces were presented upright vs. inverted (Yovel and Kanwisher, 2005). In accordance with these results, an fMRI study employing face morphing reported that activity changes in the FG, but not the STS, tracked perceptual, rather than physical, stimulus changes (Rotshtein et al., 2005). Indications of a prominent role of the FG in the perception of faces also come from a study that used Rubin’s vase ambiguous images and reported that face percepts correspond to higher activity in the FG as compared to vase percepts (Hasson et al., 2001), in spite of only minimal and peripheral stimulus changes in the two conditions. Even more direct evidence for a causal involvement of the FG in the generation of face perception comes from intracranial electrical stimulation studies that reported distortions of face perception with electrical stimulation of face selective regions of the fusiform gyrus (Puce et al., 1999; Parvizi et al., 2012; Rangarajan et al., 2014).

In spite of the lack of discriminability between upright and inverted faces in the lateral cortex, and the greater diversity observed across decoding analyses in the set of lateral electrodes, our current results cannot be interpreted as evidence of a less critical involvement of the lateral cortex in the generation of conscious face perception.

In fact, it is possible that specific regions of the lateral temporal cortex (in particular, the face selective STS) also play an important role in the generation of conscious face perception. For example, it is conceivable that STS loci contribute to aspects of face perception that are invariant to orientation. It is also possible that the information about face orientation is present in the STS, but could not be resolved by our analysis due to the coarseness of ECoG recordings, which lump the contribution of large neuronal populations. In fact, informative neuronal responses can go undetected when recorded as a mass signal, especially when informative neurons are sparse and/or weakly clustered (Dubois et al., 2015) . However, our results pose a constraint on the relative roles of ventral and temporal loci in the generation of conscious face perception.

While it is now established that the FG and the STS respond more vigorously to faces than to most other visual stimuli, the strict face-specificity of these regions is still being debated (see, for example, (Yue et al., 2011; Bilalic et al., 2011; Joseph and Gathers, 2002; Caldara et al., 2006; Haist et al., 2010; Shultz and McCarthy, 2012)). Hence, it is possible that the results we reported generalize to other, non face stimuli, especially those that are perceived as a holistic gestalt, possibly as a result of repeated exposure and hence development of configural internal templates (Diamond and Carey, 1986; Gauthier et al., 2014).

### Heterogeneity across subjects

It is important to note that our subject population is heterogeneous, especially with respect to the behavioral performance and the neurophysiological markers of conscious visibility in the masked paradigm (Fig. 2 and S7). While all subjects present at least some electrodes that discriminate visibility (v decoding, collapsing across contrast levels), the great majority of *v@c*_thr_ or *v@c*_H_ discriminant electrodes are observed in a single subject (subject 153).

Anatomical differences in electrode location across subjects are likely to underlie some of the observed diversity. In particular, only subjects 153 and 154 present electrodes in the right fusiform gyrus, which is known from several lesion, neuroimaging and electrical stimulation studies to be more markedly involved in face processing than its left homologue (e.g., (Landis et al., 1986; De Renzi et al., 1994; Tsuchiya et al., 2008; Rangarajan et al., 2014)). In addition, face-specific responses are known to be produced in small regions of the fusiform gyri, which vary in location between individuals (Allison et al., 1994; Frost and Goebel, 2012). However, these factors are unlikely to be the only source of heterogeneity. In fact, heterogeneity across subjects when decoding stimulus category in the unmasked paradigm, or when decoding *c*_3_ *vs. blank* in the masked paradigm, is much smaller than when decoding subjective phenomenology in the masked paradigm (Fig. S7 and S8). In particular, all subjects display at least one electrode in both ventral and lateral regions that is significant at p<0.001 when decoding upright face vs. other categories. Among the three subjects that were presented with inverted faces, all of them present several ventral electrodes that discriminate upright vs. inverted faces at high levels of significance (p<0.001), indicating that all of them had electrodes in regions that are highly face selective. In the masked paradigm, when considering a discrimination that is specified in terms of physical properties of stimuli such as *c*_3_ *vs. blank,* every subject displayed at least one electrode that is significant at p<0.001 in both ventral and lateral regions. However, heterogeneity across subjects increases for discriminations that are specified by subjective factors, with only subjects 153 and 168 presenting ventral electrodes that discriminate *v* at p<0.001. Heterogeneity is even higher when considering discriminations that are determined by purely subjective factors, with subject 153 presenting 9 (13) ventral electrodes that are significant at p<0.001 for decoding *v@c*_thr_ (*v@c*_H_), as opposed to none (1) for the other subjects. This suggests that at least some of the observed heterogeneity is likely to be due to different levels of task comprehension and execution, possibly resulting in visibility ratings being a poor indicator of subjective state for some subject.

Future studies could assess the extent to which intersubject variability can be accounted for by alignment of functionally corresponding brain areas across subjects. For example, one could perform preoperative fMRI recordings during a passive protocol (e.g. natural movie viewing) and use functional alignment methods to project electrode locations onto a standard brain (Sabuncu et al., 2010; Haxby et al., 2011; Conroy et al., 2013; Frost and Goebel, 2013). While functionally-based alignment has been shown to greatly reduce intersubject variability during passive tasks (Haxby et al., 2011), the extent to which functional alignment methods can account for intersubject variability in more complex and demanding cognitive tasks is an open question.

Our limited subject population does not enable us to draw definite conclusions on the origin of the observed heterogeneity. However, it is worth noting that the degree of metacognition across subjects is also variable, with 2/5 subjects (subjects 168 and 178) exhibiting negative meta-d’ at the highest contrast value tested (Fig 2C), which corresponds to higher visibility rating resulting in higher probability of incorrect response. Also, 3/5 subjects (those we just mentioned with the addition of subject 154) show considerably less than perfect objective performance (91%, 87% and 72%, for subject 154, 168 and 178) at the highest visibility rating, which corresponds to perfect visibility. Remarkably, these three subjects are the only ones that exhibit electrodes with significant *c@v* decoding (in particular, *c@v*_4_, Fig. S5), that is, with responses that discriminate face intervals with different contrast values that resulted in the same visibility rating. The correspondence between overconfident behavioral performance on the one hand, and significant *c@v*_4_ decoding on the other, suggests that these subjects might have responded with the highest visibility rating even if their perception of the face was not completely clear. As a consequence, in those trials where the highest visibility rating was reported, different contrast values would have resulted in different perceptual outcomes for these subjects. It is also worth noting that the most face selective electrodes in subject 154 were only recorded in a single session of the CFS paradigm, which did not result in enough trials for meaningful estimation of *v@c*_t_hr and *v@c*_H_ decoding accuracies.

While heterogeneity in behavioral performance and metacognition has also been reported in healthy subject populations and has been related to neuroanatomical metrics describing the local structure of grey and white matter (Fleming et al., 2010; Kanai and Rees, 2011) and to dopamine signalling (Van Opstal et al., 2014), the relationship between objective performance and metacognition on the one hand, and electrophysiological data on the other, has received less attention. It is reasonable to suppose that the heterogeneity among epilepsy patients undergoing pre-surgical monitoring might be even higher than among healthy subjects, due to variable degrees of fatigue, sleep deprivation, motivation to perform the task, possible cognitive deficits associated with long-term severe epilepsy, and possibly lingering effects from anaesthesia and/or from surgical pain. Hence, cognitive neuroscience research in implanted epilepsy subjects constitutes a precious opportunity for levering the excellent signal-to-noise ratio and high temporal and spatial resolution of intracranial recordings towards elucidating the relationship between cognitive and electrophysiological heterogeneity.

### Comparison between CFS and other masking techniques

In this study, we presented images partially masked by CFS to highlight neuronal markers that differentiate seen versus unseen trials in the absence of any changes in sensory inputs. Several other techniques allow the presentation of images at perceptual threshold, including masking and crowding. These techniques rely on different properties of the visual system and exhibit different characteristics (e.g. (Izatt et al., 2014; Faivre et al., 2014; Faivre et al., 2012; Peremen and Lamy, 2014; Kaunitz et al., 2014; Fogelson et al., 2014; Almeida et al., 2008; Tsuchiya et al., 2006), see (Kim and Blake, 2005; Macknik, 2006; Kouider and Dehaene, 2007) for reviews), hence it is possible that slightly different results could have been obtained if a technique other than CFS was used to partially mask face images.

In order to yield a deeper understanding of conscious visual perception, it is important to assess the neuronal markers of conscious visual experiences that are independent from the specific experimental paradigm employed, and those that are specific to a particular class of masking techniques. While the latter can increase our understanding of the physiological mechanisms underlying perceptual suppression (and the breaking thereof) in specific experimental paradigms (e.g., dichoptic versus monoptic masking), only the former can be considered as putative NCC-core. Many recent studies investigated the differences and similarities between masking paradigms (Izatt et al., 2014; Faivre et al., 2014; Faivre et al., 2012; Peremen and Lamy, 2014; Kaunitz et al., 2014; Fogelson et al., 2014); however, most of them employed purely psychophysical measures (Izatt et al., 2014; Faivre et al., 2012; Peremen and Lamy, 2014; Kaunitz et al., 2014), or non-invasive electrophysiological measures with poor spatial and temporal resolution (Fogelson et al., 2014). Hence, the comparison between neuronal markers of conscious visual perception under different masking paradigms is an important topic for future research, and one that can greatly benefit from the high temporal and spatial specificities of intracranial recordings.

### Distilling NCC-core - caveats and potential confounds

It is important to recognize that the combination of masked and unmasked paradigms can greatly contribute towards a more accurate assessment of the core neuronal correlates of conscious visual perception, but also presents some caveats. For example, some electrodes can be upright face discriminant and visibility discriminant, but still not NCC-core. For example, areas related to memory formation of faces would be both face discriminant and visibility discriminant, but not NCC-core, in the sense that a hypothetical patient with a localized lesion in that area might still be able to experience faces consciously, albeit incapable of creating long-lasting memories of face identities (Postle, 2009).

More generally, an electrophysiological feature that distinguishes both high vs. low visibility trials in the masked paradigm and upright faces vs. other stimuli in the unmasked paradigm does not necessarily constitute an NCC-core marker, since it could represent a correlate of the same confound in both scenarios. For example, one electrode from the temporal pole of subject 178 exhibited significant A’ both in a CFS visibility decoding analysis and in an unmasked face decoding analysis (Fig. S6). While face selective effects have been reported also in anterior temporal and frontal regions, especially in the anterior ventral temporal cortex, inferior frontal gyri, frontal operculum and lateral prefrontal cortex (Allison et al., 1999; Avidan et al., 2005; Chan et al., 2006; Kriegeskorte et al., 2007; Tsao et al., 2008; Engell and Haxby, 2007; Said et al., 2010; James et al., 2013), the anatomical location of the face discriminant electrode in the temporal pole of subject 178 is also compatible with discriminant activity in this region being due to eye movements towards the eyes of the target face both in high visibility CFS trials and in unmasked trials containing a face (Jerbi et al., 2009; Kovach et al., 2011). Future work can disambiguate the influence of eye movement confounds by simultaneous recordings of eye movements.

In this study we focused on local neuronal responses. However, it has been suggested that conscious awareness relies on functional coupling across neuronal populations (Engel et al., 1999; Thompson and Varela, 2001; Melloni et al., 2007; Godwin et al., 2015); more specifically, on a network of irreducible causal interactions across neuronal populations (Seth et al., 2011; Oizumi et al., 2014; Tononi and Koch, 2015; Haun et al., 2016). The development of improved methods for the assessment of causal interactions in neuronal data (Oizumi et al., 2016), as well as advances in brain imaging and large-scale neural recording and stimulation techniques (Duyn, 2012; Ahrens et al., 2013; Shobe et al., 2015; Buzsaki et al., 2015; Yang et al., 2016), are expected to greatly foster the investigation of multidimensional patterns of neuronal interactions as putative neuronal correlates of consciousness.

## Conclusion

We have combined a set of decoding analyses of either physical properties of stimuli or subjective phenomenology in a masked task, with a set of decoding analyses of stimulus category in an unmasked task. Our results in the masked task show that subjective phenomenology is better decodable than physical properties of stimuli (such as contrast) in the high level visual areas considered in this study, i.e. the ventral and lateral sides of the temporal lobe.

While we were able to decode subjective visibility in the masked task with high accuracy in both ventral and lateral loci, the inclusion of a stimulus category decoding analysis in the unmasked task revealed an important difference between the two loci: while both ventral and lateral areas discriminate upright faces from other stimuli, only ventral areas discriminate upright from inverted faces.

Our results suggests a critical role for ventral brain areas, and in particular for the fusiform gyrus, in the conscious configural perception of faces and possibly other objects that are perceived holistically. More generally, this work points towards a promising direction in consciousness research based on the combination of similar protocols tailored to expose specific aspects of the conscious visual experience.

## Acknowledgments

We thank all subjects for their participation in the study, the medical team at University of Iowa Hospital for their assistance, Haiming Chen for his technical assistance. We thank Andrew Haun for generating Figure 1. NT is supported by Australian Research Council (ARC) Future Fellowship (FT120100619) and Discovery Project (DP130100194). CKK, HK, HO, and MAH were supported by NIH grants R01 DC004290 and UL1RR024979.

## Competing Interests

NT collaborates with Dr Ryota Kanai, CEO of a venture company, called Araya Brain Imaging (araya.org) based in Tokyo, Japan. In October 2015, Araya received a grant from Japan Science and Technology (JST; approximately 300 thousand yen for 5 years), which aims to potentially commercial projects related to artificial consciousness. NT is part of this grant and he is supposed to host several students/research assistants/postdocs hired at Araya over the next 5 years. However, his institute (Monash University) does not receive any research income directly from JST. NT does not hold stocks or shares of Araya. NT is not and will not be paid by Araya. As the collaboration has barely started, no patent application has been submitted. Some of NT’s travels are covered by the JST through Araya. Please note that the research reported here is unrelated with the grant from JST, which just started on November 2015.

## Author Contributions

Conceived the idea of the study: FB NT. Conceived and designed the experiments: HK CKK MAH RA NT. Performed neurosurgery: HK MAH. Collected the data: HK CKK NT. Conducted preliminary analyses including anatomical localization of electrodes: HK HO NT. Analyzed the data: FB JvK NT. Wrote the manuscript: FB NT. Commented critically on the manuscript: FB JvK HK CKK HO MAH RA NT.

## Supplementary Figures

**Figure S1:**
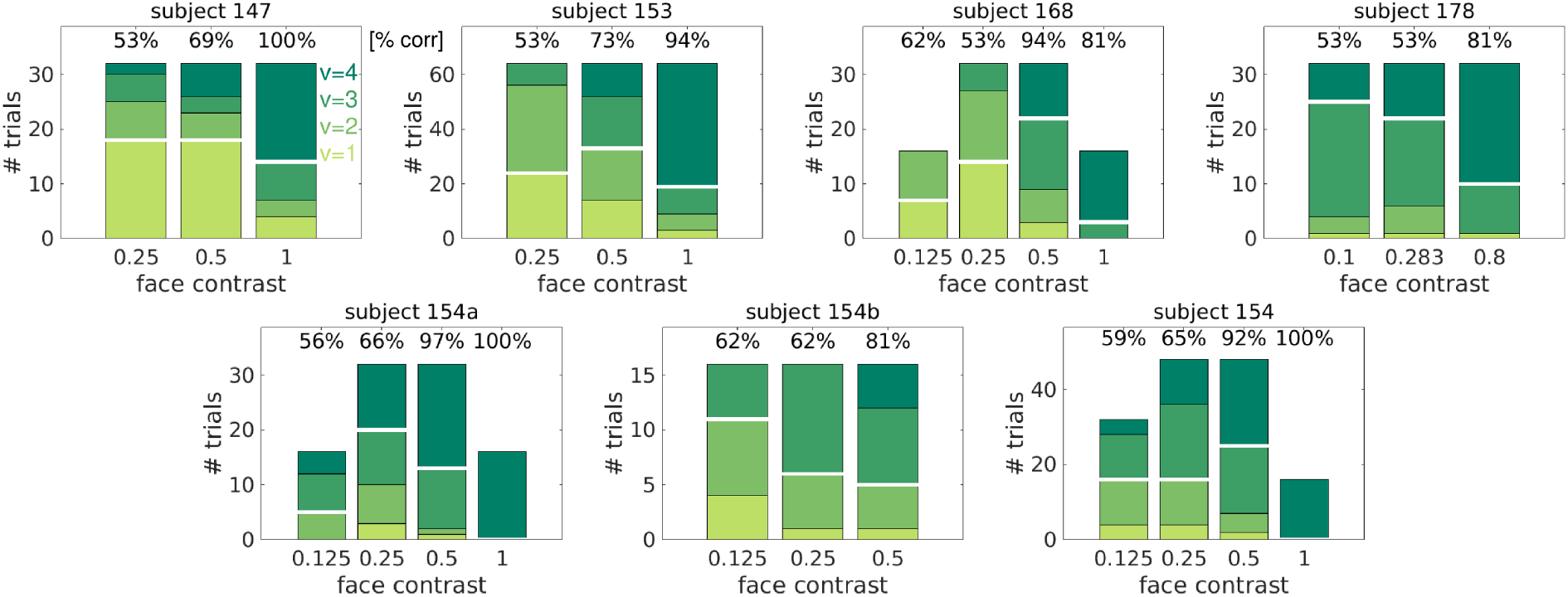
CFS behavioral results for each subject. The stacked bar plots show, for each contrast value, the number of trials that have been assigned to each visibility rating. Color code as in Fig. 2E,F. Numbers on top of each bar show the objective performance (percentage of correct trials over all trials) for each contrast value. In the case of subject 154, the set of recorded electrodes differ across sessions: “154a” and “154b” indicate the sets of electrodes that have been recorded in only a subset of the sessions, while “154” indicates the set of electrodes that has been recorded in every session.

**Figure S2.**
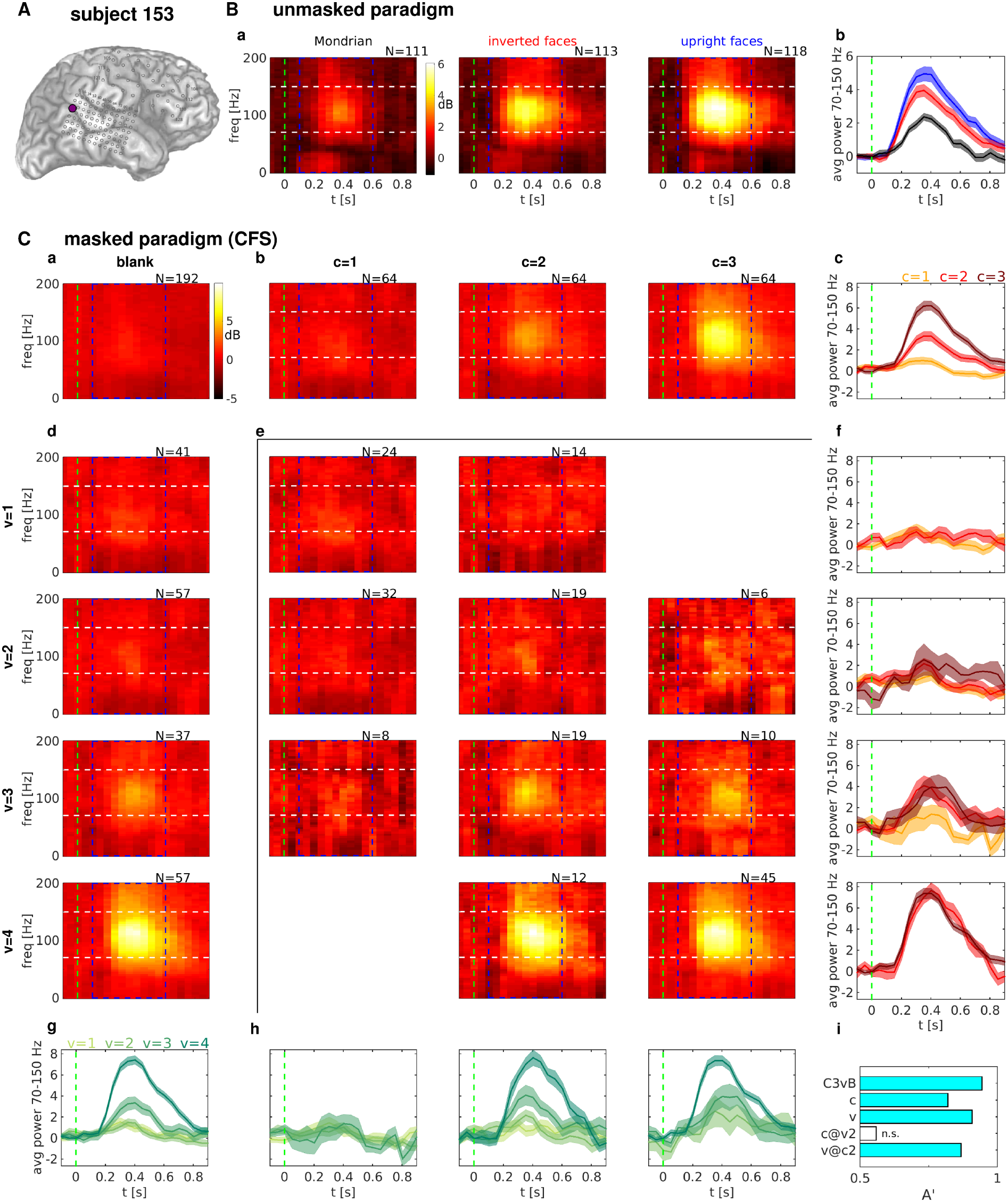
Spectral power responses in the unmasked and CFS experiment for an example lateral electrode. Spectral power responses in the unmasked and CFS experiment for an example electrode. Format as in Figure 3.

**Figure S3.**
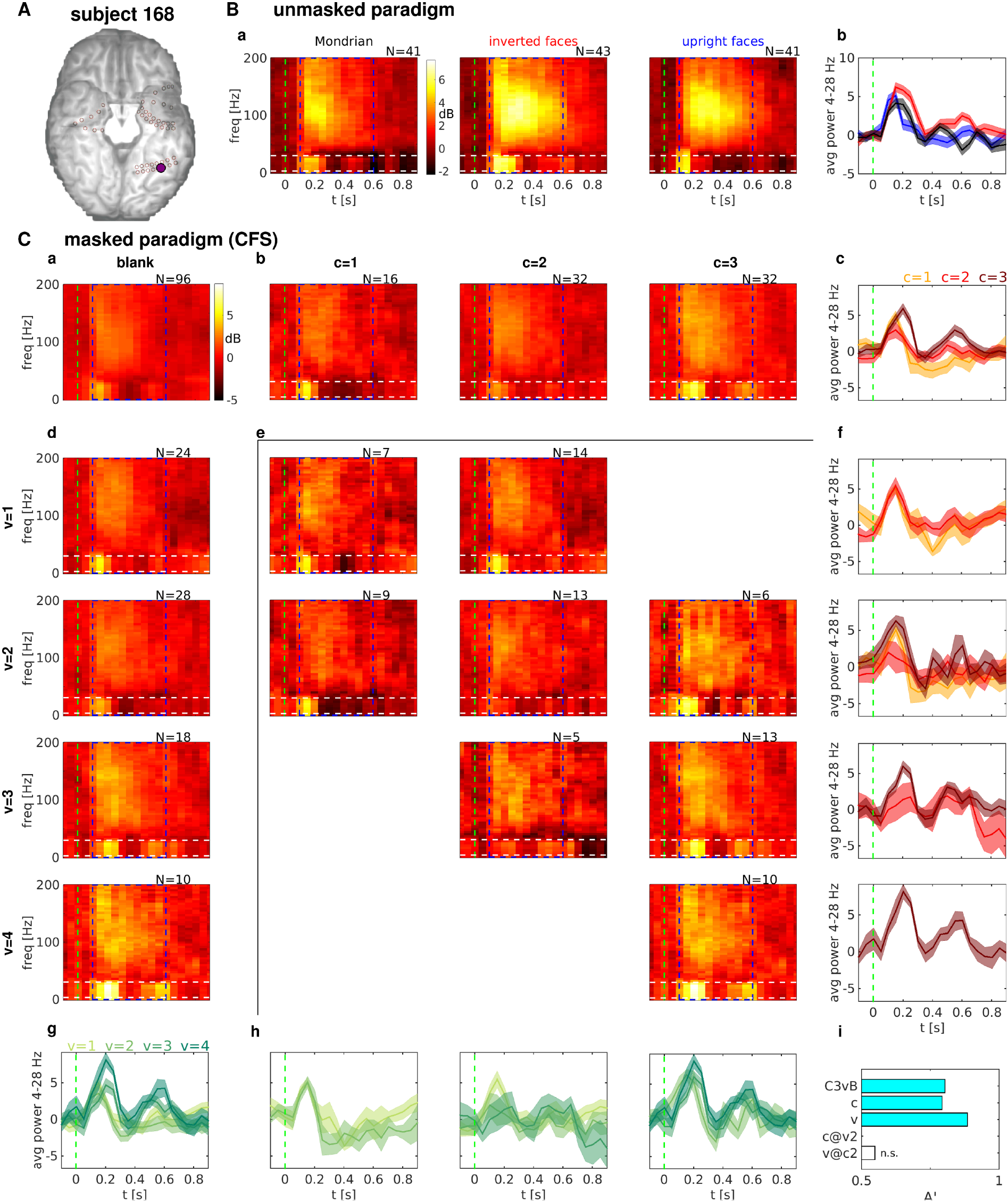
Spectral power responses in the unmasked and CFS experiment for an example ventral electrode. Spectral power responses in the unmasked and CFS experiment for an example electrode. Format as in Figure 3. The trials with face contrast value *c*=4 have not been considered for this figure.

**Figure S4.**
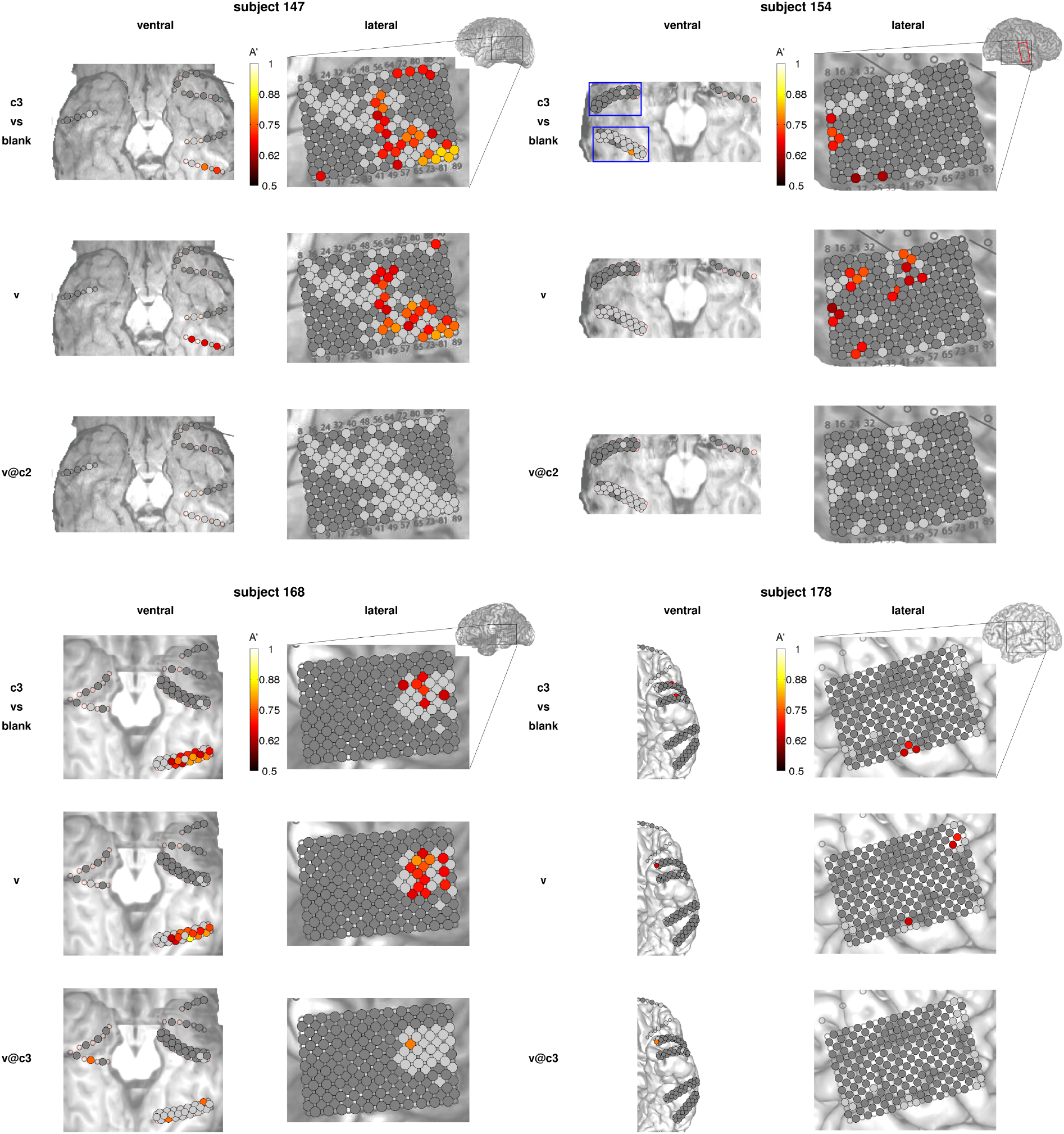
Decoding accuracies in the masked paradigm. Decoding accuracies A’ for each electrode are shown color-coded on the ventral and lateral brain images for the remaining four subjects. Same format as in Fig. 4A. For subject 154, different electrodes were recorded on different sessions: the red rectangle in the lateral temporal area indicates the electrodes that have been recorded in the sessions corresponding to “154a” only, and the blue rectangles in the ventral temporal area indicate the electrodes that have been recorded in the sessions corresponding to “154b” only. The other electrodes have been recorded in every session.

**Figure S5.**
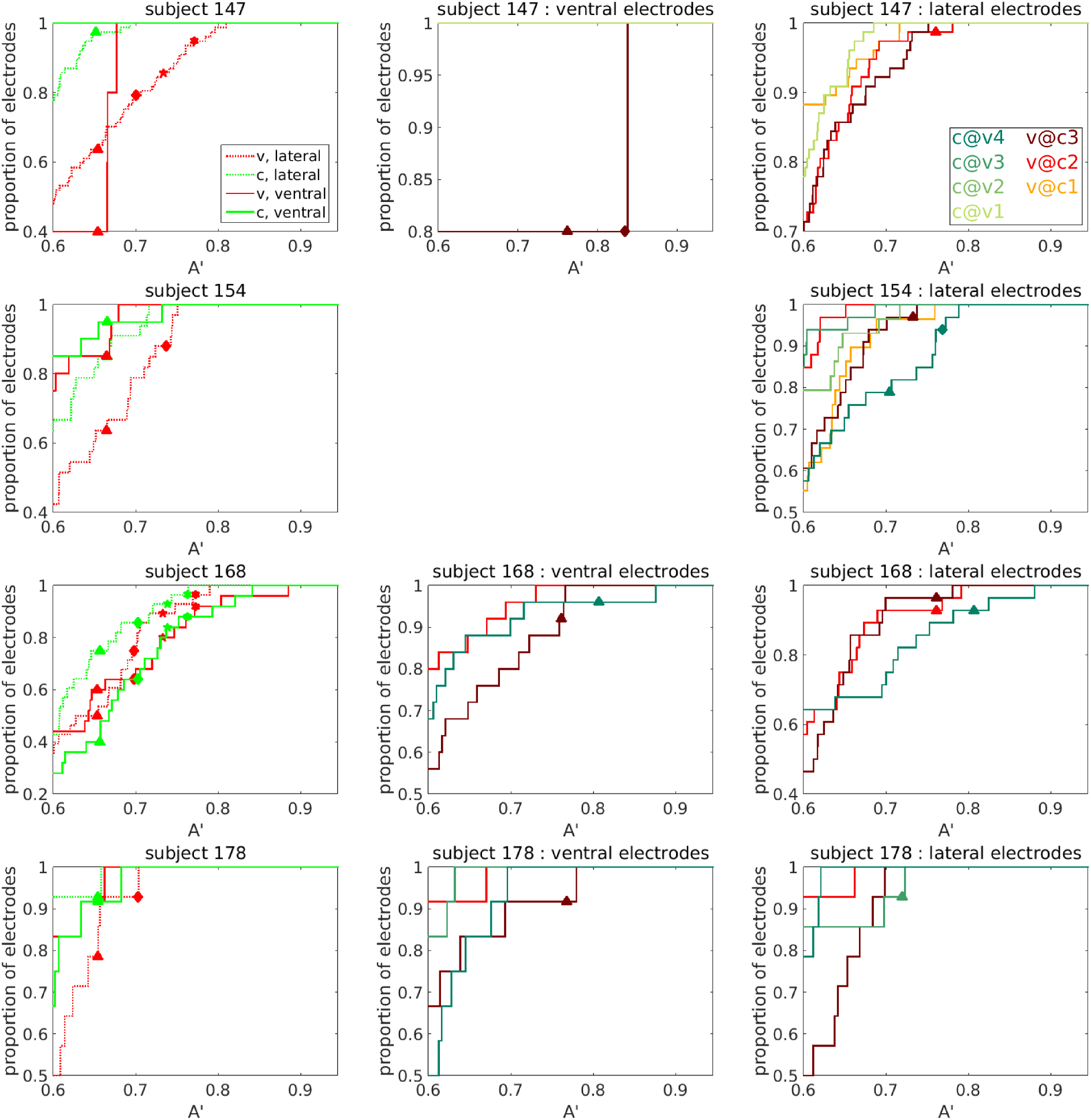
Cumulative probability density functions of decoding accuracy over the populations of ventral and lateral electrodes. Same format as in Fig. 6B, for the remaining subjects. Only results from decoding analyses with at least 10 trials in the least populated class are shown. For subject 154, the numbers of electrodes considered are different for each decoding analysis, as indicated in Fig. S1 and S4. For this subject, none of the face-responsive electrodes located in the ventral cortex have enough trials to compute *v@c* or *c@v* decoding, hence the corresponding panel is not shown.

**Figure S6.**
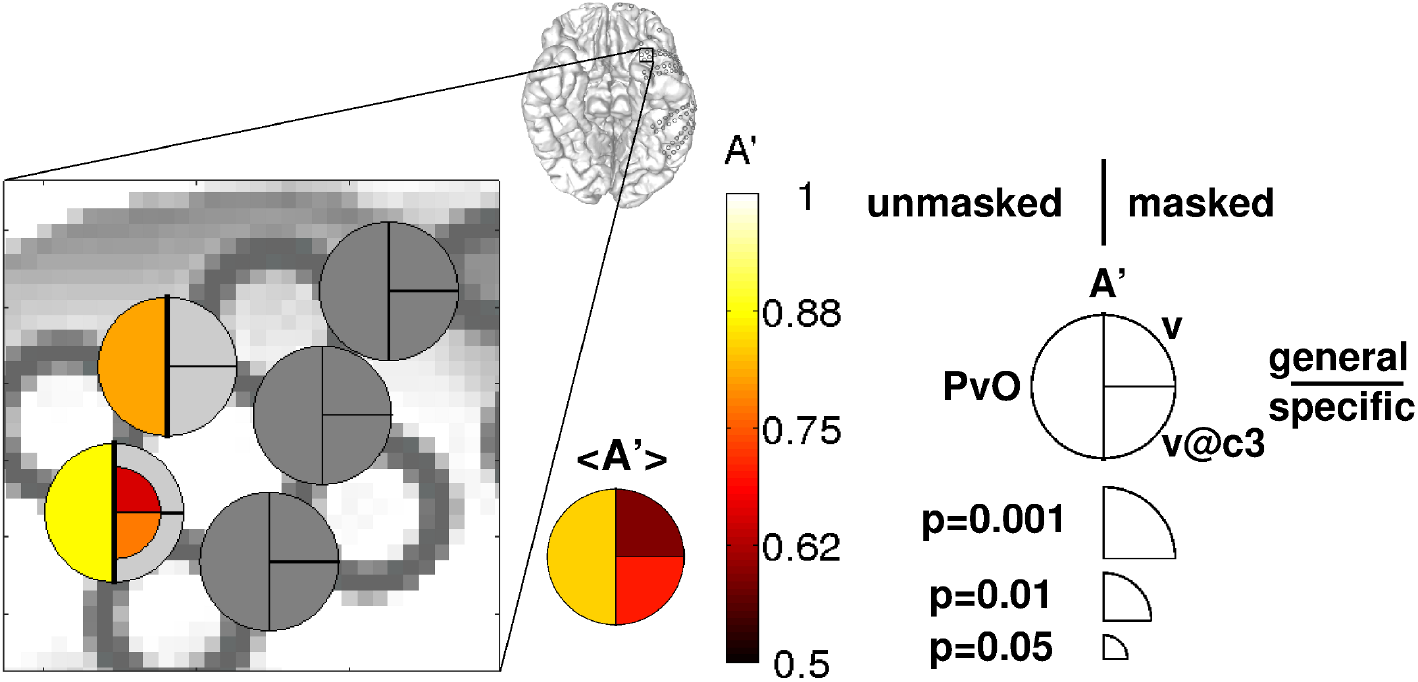
Comparison between masked and unmasked presentations: example of a visibility discriminant and face discriminant electrode located in the temporal pole. Same format as in Fig. 7A, for a set of face discriminant electrodes from the temporal pole of subject 178.

**Figure S7.**
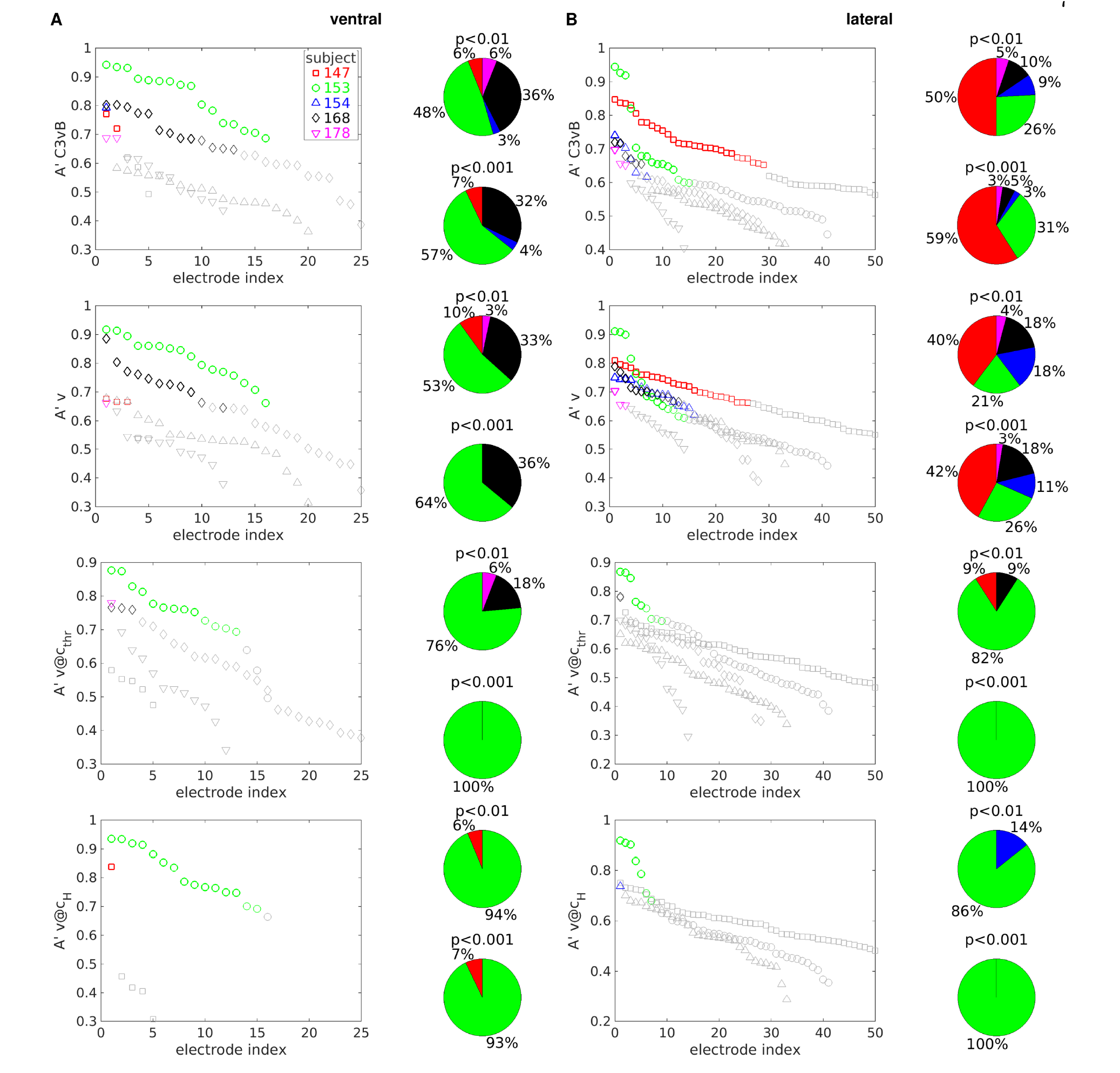
Heterogeneity of decoding results across subjects in the masked paradigm. (A) Decoding accuracies A’ for each subject and each ventral electrode are shown in descending order. Thick (thin) colored symbols indicate significant decoding accuracy at p<0.001 (p<0.01), gray symbols indicate non-significant decoding accuracy (p>0.01). Only the 50 electrodes with the highest A’ for each subject and brain region are shown. Colors as in Fig. 2A-D, symbols as in Fig. 5A. Pie charts show the proportion of significant (p<0.01, top; p<0.001, bottom) lateral electrodes contributed by each subject for each decoding analyses. From top to bottom: *c*_3_ *vs. blank*, *v*, *v*@*c*_thr_, *v@c*_H_. (B) As in A, for lateral electrodes.

**Figure S8.**
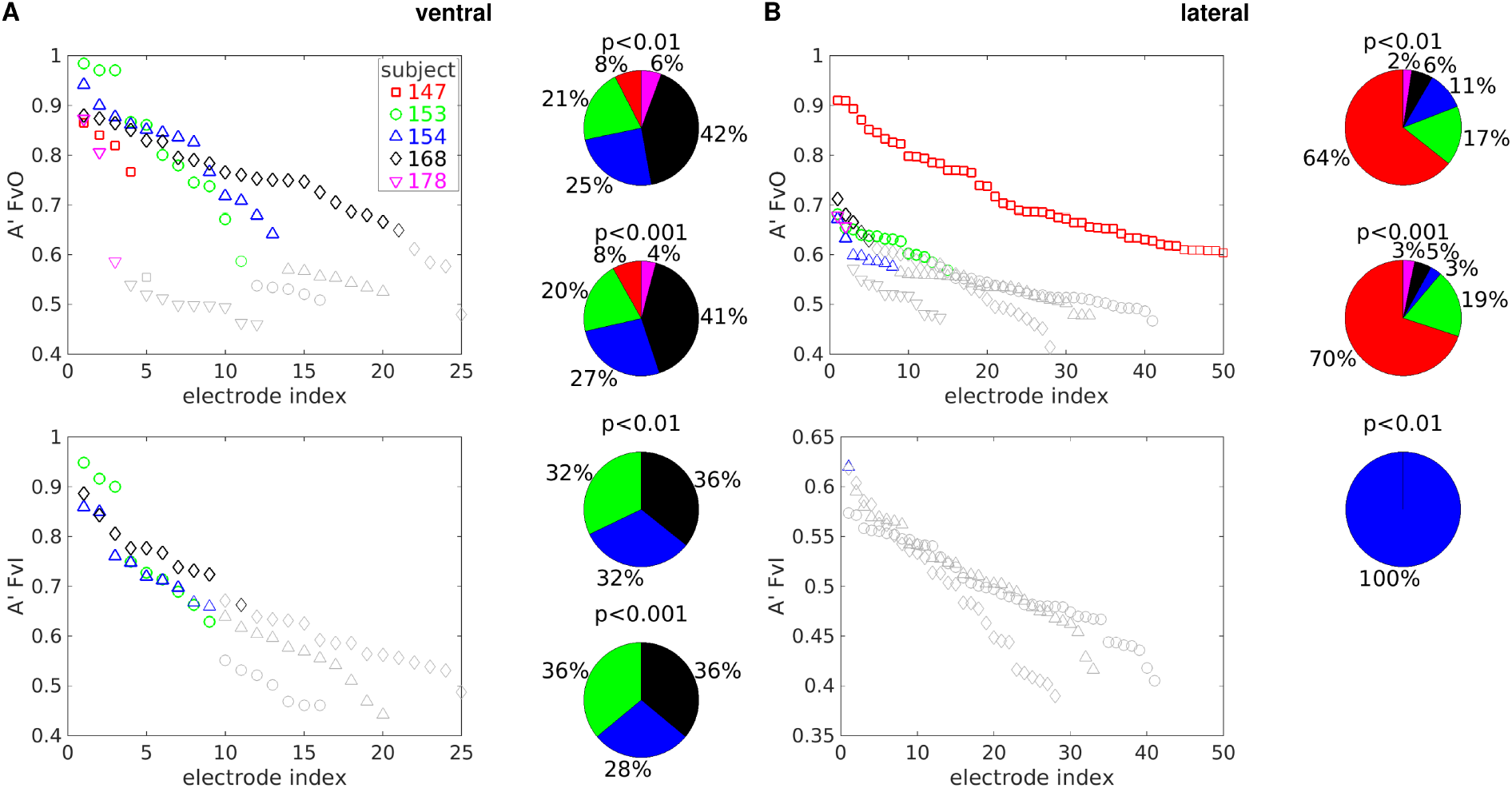
Heterogeneity of decoding results across subjects in the unmasked paradigm. (A) Decoding accuracies A’ for each subject and each ventral electrode are shown in descending order. Thick (thin) colored symbols indicate significant decoding accuracy at p<0.001 (p<0.01), gray symbols indicate non-significant decoding accuracy (p>0.01). Only the 50 electrodes with the highest A’ for each subject and brain region are shown. Colors as in Fig. 2A-D, symbols as in Fig. 5A. Pie charts show the proportion of significant (p<0.01, top; p<0.001, bottom) lateral electrodes contributed by each subject for each decoding analyses. From top to bottom: upright face vs. other categories, upright vs. inverted face. (B) As in A, for lateral electrodes.

